# Multi-Level Regulation in RNA-Protein Hybrid Incoherent Feedforward Loop Circuits for Tunable Pulse Dynamics in *Escherichia coli*

**DOI:** 10.1101/2025.06.20.660716

**Authors:** Seongho Hong, Syeda Simra Shoaib, Mathias Foo, Xun Tang, Jongmin Kim

**Author notes:** **Corresponding Authors** Mathias Foo - School of Engineering, University of Warwick, Coventry CV4 7AL, UK;, Xun Tang - Cain Department of Chemical Engineering, Louisiana State University, Baton Rouge, Louisiana 70803, USA;, Jongmin Kim - Department of Life Sciences, Pohang University of Science and Technology, Pohang 37673, Republic of Korea. These authors contributed equally (Seongho Hong, Syeda Simra Shoaib).

## Abstract

Regulating gene expression with precision is essential for cellular engineering and biosensing applications, where rapid, programmable, and sensitive control is desired. Current approaches to regulatory circuit design often rely on control at a single regulatory level, primarily the transcriptional level, thereby limiting the capability of fine-tuning the regulatory dynamics in response to complex stimuli. To address this challenge, we developed four novel RNA-protein hybrid type-1 incoherent feed-forward loop (I1-FFL) circuits in *Escherichia coli* that integrate transcriptional and translational regulators to achieve multi-level control of gene expression. These hybrid circuits leverage the modularity and rapid dynamics of RNA-based activators alongside the versatile inhibition capabilities of the protein-based repressors, to endow tunable pulse dynamics through engineered delays that act as transient repressor decoys. By repurposing synthetic RNA regulators at multiple regulatory levels together with aptamer and RNA-binding proteins, we demonstrate previously unexplored circuits with tunable dynamics. Complementary simulation results highlighted the importance of the engineered delays in achieving tunable pulse dynamics in these circuits. Integrating modeling insights with experimental validation, we demonstrated the flexibility of designing the RNA-protein hybrid I1-FFL circuits, as well as the tunability of their dynamics, highlighting their suitability for applications in environmental monitoring, metabolic engineering, and other <u>engineered biological systems, where precise temporal control and adaptable gene regulation are desired.</u>

---

Synthetic biology has become a powerful approach for engineering artificial decision-making circuits within living organisms^1^, enabling precise and programmable regulation of gene expression^2^. Over the past few decades, remarkable progress has been made in constructing synthetic gene circuits, such as oscillators^3^, bistable switches^4^, and arithmetic circuits^5^, to showcase the unprecedented programmability and potential applications of synthetic biology. These advancements have paved the way for novel applications of synthetic gene circuits in domains that demand highly sensitive detection and control^6^, from biosensor development to chemical production and immunomodulation-based cell therapies^7, 8^.

RNA-based regulators have gained prominence due to their exceptional programmability and rapid response properties^9, 10^, making them ideal components in biosensing technologies^11^. Recent breakthroughs have introduced *de novo*-designed synthetic RNA regulators, such as small transcription activating RNAs (STAR)^12^ and toehold switches (THS)^13^, enabling precise regulations at the transcriptional (TX) and translational (TL) levels of gene expressions. These versatile RNA tools offer extensive libraries, dynamic ranges, and modularity^14^, making them ideal components for constructing complex synthetic gene circuits^15, 16^ that regulate gene expression based on environmental cues.

The incoherent type-1 feed-forward loop (I1-FFL) network motif^17^ is a widely studied natural network in synthetic and systems biology. Consisting of three interacting nodes (X, Y, and Z), the I1FFL circuit can generate a pulse in node Z gene expression through two incoherent regulatory paths. This phenomenon is associated with diverse biological processes, such as stress response^18^, signaling^19^, and development^20^. Recent studies have demonstrated the utility of I1-FFLs in biosensing, particularly in environments that demand temporal adaptation^21^ and efficient signal detection^22^. I1-FFLs have been shown to facilitate effective long-distance communication in microbial consortia^23^, allowing coordinated signal propagation across spatially distributed populations. In mammalian cells, I1-FFLs help alleviate gene expression burden by optimizing resource use^24^ and enhancing signal fidelity^25^, and enabling precise metabolic control for improved circuit performance^26^. Moreover, their ability to detect fold changes rather than absolute concentrations makes them ideal for decoding and responding to oscillatory signals^27^. Additionally, I1-FFLs offer temporal adaptation, allowing systems to return to baseline after prolonged stimuli, and have proven effective in bacterial systems for robust adaptive control in fluctuating environments^28^.

Synthetic I1-FFL circuits often employ TX regulators^29-31^ such as transcription factors to modulate the gene expression by targeting the promoter regions. While effective, targeting relatively short promoter regions using transcription factors could limit the design flexibility to achieve tunable dynamics. In this work, we present a novel approach to establish multi-level inhibition pathways, by incorporating RNA-binding proteins (RBPs)^32^ in tandem with aptamers. Unlike TX regulators constrained by promoter configurations, the RBP-based inhibition with aptamers offers a more flexible and modular alternative for controlling the gene expression activity at the TL level^33^. For instance, RBP can be clustered to a far greater extent than transcription factors by using aptamer arrays^34^, and thus, potentially enhancing the inhibition efficiency. Furthermore, combining different types of aptamers, such as MS2 and Qβ^35^, could allow the design of complex regulatory logic, and enable combinatorial configurations for constructing complex logic circuits.

In our previous study, we explored several RNA-only and RNA-protein hybrid I1-FFL circuits in *Escherichia coli* (*E. coli*)^36, 37^. We found that while RNA-only I1-FFL circuits demonstrated rapid kinetics due to RNA’s faster production and degradation rates than proteins, they lacked significant temporal separation between the activation and the inhibition pathways for a pronounced pulse-like profile in the target gene expression. On the other hand, the integration of protein-based regulatory components in the RNA-protein hybrid I1-FFL design could introduce a sufficient timescale difference between the two incoherent regulation pathways to yield pulse-like dynamics. Based on these findings, here we further explore the design and tunability of four novel RNA-protein hybrid I1-FFL circuits, evaluating their capability of generating a pulsed expression in the target gene, adding to the pool of genetic pulse generators.

In this study, we developed a novel framework for multi-level gene regulation by constructing four RNA-protein hybrid I1-FFL circuits in *E. coli*. By integrating combinations of TX and TL regulators, including STAR, THS, TetR, and RBP, we systematically investigated how multi-level regulation enables precise control over circuit dynamics. This integration bridges the rapid activation capabilities of RNA-based regulators with the tunable repression strengths of protein-based repressors, overcoming the limitations of single-level regulation in achieving dynamic pulse generation and temporal tuning. Additionally, we developed mathematical models to analyze and optimize the circuit behavior, providing a quantitative framework for designing adaptable gene regulatory networks. These hybrid circuits not only expand the versatility and tunability of I1-FFL circuits, they also serve as a flexible platform for biosensing applications such as environmental monitoring, therapeutic gene expression, and metabolic engineering, where precise temporal regulation and robust adaptability are required.

## RESULTS

### Design of RNA-Protein Hybrid I1-FFL Circuits for Multi-Level Regulation

To systematically investigate the effects of multi-level regulation on gene expression dynamics, we designed and constructed four distinct RNA-protein hybrid I1-FFL circuits, with each featuring a different combination of TX and TL activators and inhibitors. These circuits were strategically chosen to explore how TX and TL regulatory mechanisms contribute to pulse generation and tunability. Each circuit was named based on the regulation type at each node for simplicity (Figure 1A). For example, the TL-TX-TL circuit features a TL activation of node Y by node X, a TX activation of node Z by node X, and a TL repression of node Z by node Y.

**Figure 1.**
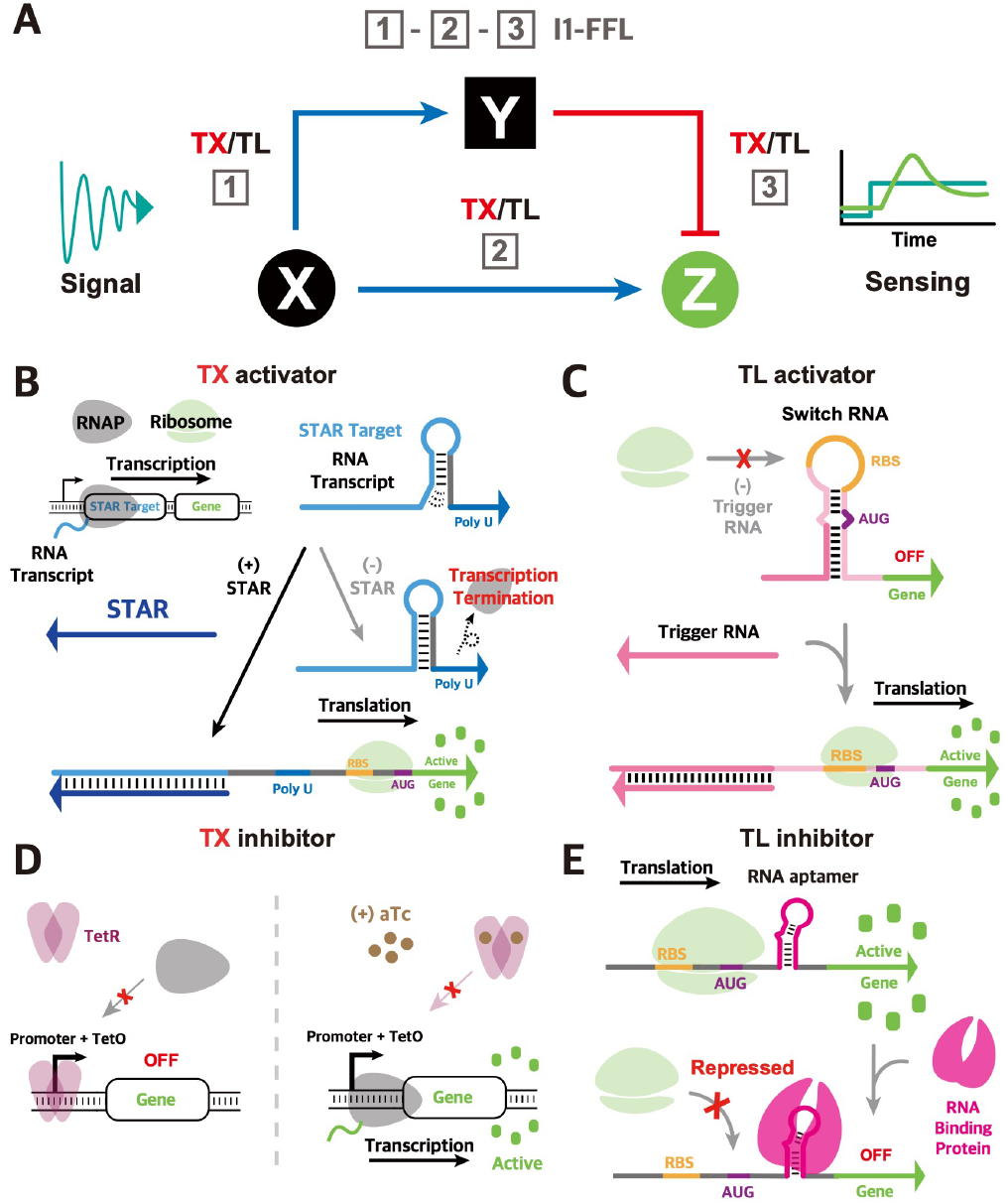
Schematics of Regulatory Components in I1-FFL Circuits. (A) Overview of I1-FFL architecture with a combination of TX and TL controls. (B) STAR activates transcription by unwinding terminator structures, initiating downstream gene expression. (C) THS enables translation by unwinding its secondary structure, exposing the ribosome binding site (RBS) for ribosome access. (D) TetR represses transcription by binding to TetO operator, where addition of aTc releases TetR from TetO, thereby modulating repression. (E) RBP inhibits translation by binding to an up- <u>stream aptamer, blocking ribosome access to the RBS.</u>

For all the designs, the input node X provides both STAR and THS triggers under the control of T7 promoter to activate the intermediate node Y and the output node Z (Figure S1). To minimize crosstalk and ensure modularity, the X, Y, and Z nodes were each cloned into different plasmids, co-transformed into the *E. coli* BL21(DE3) cells^41^, and controlled by IPTG induction. Different promoter combinations were employed to optimize circuit behavior (Figures S2, S3). Further experimental details can be found in Supplementary Information.

Each regulatory component was selected for its distinct mechanism of action. The STAR TX activation mechanism^12^ prevents transcription termination by binding to the 5′ region of the STAR target RNA, which otherwise forms a terminator structure that blocks transcription of the downstream gene. By introducing STAR RNA, the terminator structure is disrupted, enabling transcription to proceed (Figure 1B). The THS TL activation mechanism^13^ involves a trigger RNA that interacts with a translationally inhibited target gene, where the RBS and start codon are sequestered in a stable secondary structure. This structure prevents ribosome access, but upon interaction with the trigger RNA, the hairpin unwinds to expose the RBS and start codon, allowing translation to initiate (Figure 1C).

For the protein inhibitors, we employed TetR TX inhibitor and RBP TL inhibitor. The TetR TX inhibitor interferes with the entrance of RNA polymerase by binding to the operator region of pTet promoter^42^. TetR functions as a dimer with high binding affinity, conferring an efficient repression even at low concentrations. Allosteric binding of tetracycline analogs, such as aTc, could induce a conformational change in TetR to reduce its binding affinity, thereby offering opportunities to tune the target gene expression (Figure 1D). RBP, specifically coat proteins from viruses such as PP7, functions as a TL inhibitor by binding to its cognate RNA aptamer upstream of the target gene to block ribosome activity to control the translation of target RNA^35^ (Figure 1E).

As the intermediate node Y could play an important role for finetuning the circuit dynamics, we focused on the exploration of diverse tools to modulate the dynamics of node Y. To tune the expression levels at node Y, we tested different synthetic constitutive promoters, and employed different synthetic protein degradation tags^43^ to tune the lifetime of the protein inhibitors. By varying these regulatory elements, we aimed to elucidate the impact of node Y dynamics on pulse generation. Specifically, we examined the interplay of the promoter strength and degradation rate to determine how these parameters influence the system’s ability to generate pulses and maintain temporal separation between activation and repression pathways.

### Analysis of I1-FFL Circuits Using TX Inhibitor TetR as Node Y

To explore the influence of TX regulation on pulse dynamics, we analyzed the performance of two TetR-based I1-FFL circuits: TX-TL-TX and TL-TX-TX. These circuits were designed to investigate how variations in TetR production and degradation affect pulse-like gene expression, leveraging both experimental measurements and computational simulations to provide insights into the interplay between TX activation and inhibition.

In the TX-TL-TX circuit, node X produces both STAR and THS triggers to activate node Y and Z, respectively (Figure 2A). Upon activation by STAR, node Y constitutively expresses TetR protein. Node Z produces the reporter/target gene GFP, expressed by the T7 promoter, which contains both a TetO operator sequence for repression by TetR and a THS target sequence for activation by trigger from node X. Experimental measurements revealed that the pulse dynamics is tunable by varying the promoter strength for TetR expression. Using strong promoters, such as J23119, for TetR expression could lead to prominent early peaks when compared to using weak promoters, such as J23116 (Figure 2B). The simulation results indicated that the circuits peaked earlier when a stronger promoter is used for Y (Figure 2C, S4A).

**Figure 2.**
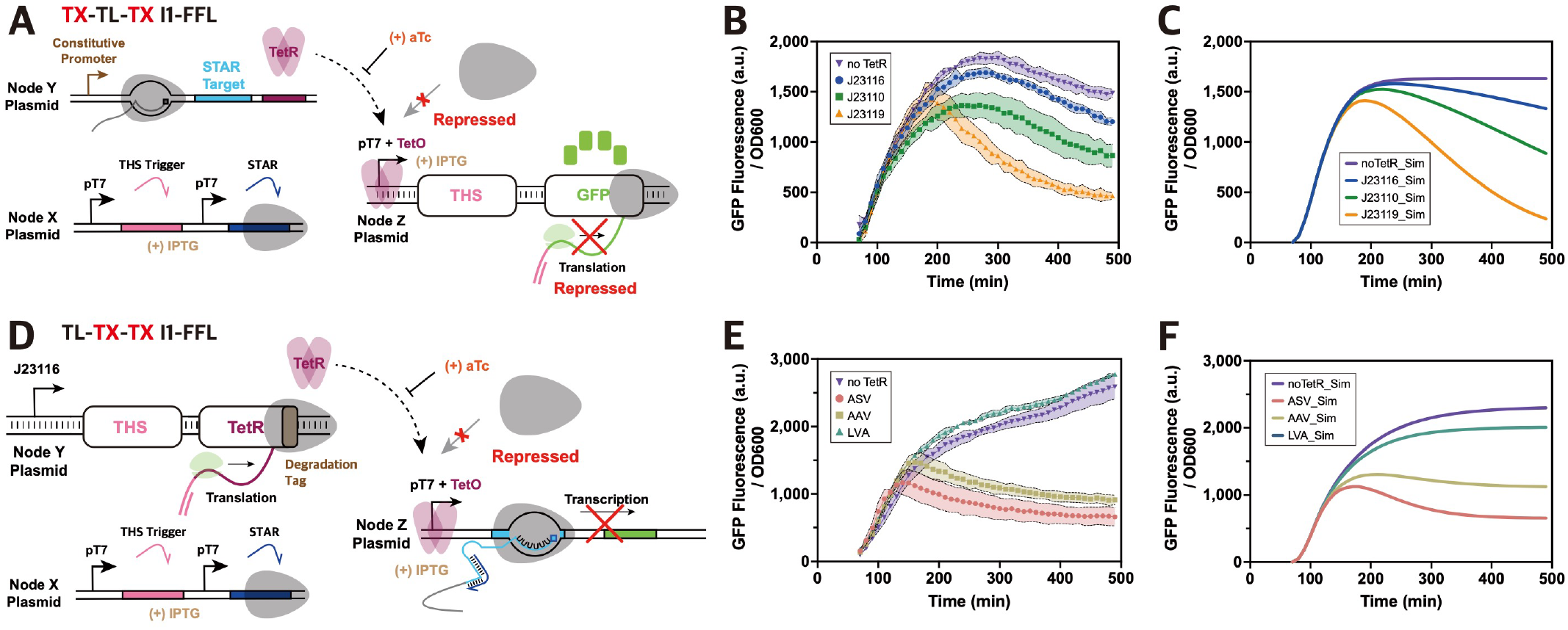
Characterization of I1-FFL Circuits Using TetR as Node Y. (A) TX-TL-TX circuit design: Node Y is activated by STAR, and node Z by the THS trigger. TetR from node Y represses node Z transcription at the TetO region, adjusted by aTc. (B) TX-TL-TX circuit experimental results: GFP expression from node Z measured over 490 min with node Y expressed using different promoters. aTc was treated at 50 ng/mL. GFP fluorescence normalized by OD600 values are plotted, where dots represent the average with shades representing standard deviation of biological triplicate measurements. (C) Simulation results for TX-TL-TX circuit model. (D) TL-TX-TX circuit design: TetR, activated by THS, represses node Z expressed by STAR. (E) TL-TX-TX circuit experimental results: GFP expression from node Z with varying TetR degradation tags. aTc was treated at 100 ng/mL. GFP fluorescence normalized by OD600 values are plotted, where dots represent the average with shades representing standard deviation of biological triplicate measurements. (F) Simulation results <u>for TL-TX-TX circuit model.</u>

In the TL-TX-TX I1-FFL circuit, an analogous design was used, except that node Y is now activated by THS, and node Z is activated by STAR (Figure 2D). A weak J23116 promoter was employed for node Y to partially address the expected high gene expression level from THS. Rather than increasing promoter strength, we aimed to modulate the associated pulse dynamics by varying TetR degradation rates. Experimentally, circuits with weaker degradation tags (AAV or ASV) for TetR produced more prominent peaks than that with a strong tag (LVA) (Figure 2E). Presumably, weaker degradation tags could ultimately lead to a sufficient accumulation of TetR over time for a strong repression with some delay, thereby resulting in prominent peaks. Model-based analysis on the relative abundance of TetR and STAR in the system revealed that, the concentration difference between TetR and STAR is more pronounced in circuits with a weaker degradation tag (ASV), as compared to circuits with a strong tag (LVA) (Figure S4). Such a high concentration difference between TetR (ASV-tagged) and STAR may have contributed to pulse generation.

Furthermore, adjusting aTc concentrations demonstrated the tunablility of TetR-based repression in both the TX-TL-TX and the TL-TX-TX circuits (Figures S5, S6). Higher aTc concentrations reduced TetR binding to TetO, delaying the repression and shifting the pulsatile GFP expression to a later time. These observations are consistent across all circuits with different promoter variants, suggesting the reliability of the aTc-based TetR repression tunability. The availability of numerous TetR variants^42^ further provides a wide range of options to use the aTc-based TetR repression mechanism, to fine-tune gene expression for precise control over genetic regulation^44^.

Together, the demonstration of pulsed gene expression in the two circuits also indicates a significant timescale difference (i.e., the condition for pulse generation^45^) between the activation and the repression pathways, suggesting that RNA-based activators act at a faster rate as compared to the protein-based inhibitor.

Characterization of PP7 Aptamer-Based TL Inhibitor. The ability to regulate gene expression at multiple checkpoints is crucial for creating complex and finely tuned genetic circuits. To expand the synthetic regulatory parts for TL regulation in dynamical circuits, we employed RBP as the TL inhibitor at node Y within the I1-FFL circuits as novel TL inhibition mechanism. PP7 coat protein (PP7), along with MS2 coat protein, is one of the most-commonly used RBP^46^, due to the detailed characterization and mechanical understanding of its functionality. The PP7 binding to its aptamer is stable enough to disrupt the ribosome entry if the aptamer is placed proximal to the translation initiation site^40^, offering opportunities for controlled TL inhibition. Thus, we aimed to explore the incorporation of aptamer in the output node Z and its impact on TL regulation.

Still, due to the structural differences for STAR and THS, we need to first elucidate the requirements for effective TL regulation in combination with STAR and THS. For the STAR design, the RBS and start codon are located downstream of the STAR target, such that the placement for PP7 aptamer is flexible with expected repression of downstream GFP translation upon activation by the STAR trigger (Figure S7A). Whereas for the THS design, the RBS and start codon are sequestered within a stable hairpin structure such that PP7 aptamer cannot immediately follow the start codon (Figure S7B).

To optimize the repression efficacy, we chose to evaluate spacer lengths between 8 and 14 nucleotides (nt) of the aptamer in relation to the start codon position (Figure S7C), since efficient repression of translation was reported if the aptamer is located within 15 nt downstream of the start codon^40^. GFP fluorescence measurements from plate reader assays revealed that spacers of 10 nt or fewer achieved over 90% repression for both STAR and THS when PP7 was present (Figure S7D,E; Tables S2,S3). The highest GFP expression level in the absence of PP7 occurred with the 9 nt spacer for STAR and with the 10 nt spacer for THS, indicating the critical role of sequence context and secondary structure in determining TL efficiency. Consequently, the final circuit designs adopted the 9 nt spacer for STAR and the 10 nt spacer for THS, ensuring optimal repression efficiency for both systems.

Analysis of I1-FFL Circuits Using RBP as Node Y. Building on our characterization of PP7 aptamer-based TL inhibition and its integration into I1-FFL circuits, we next investigated how RBP, as a TL inhibitor, influences the dynamic properties of RNA-protein hybrid I1-FFL circuits. Specifically, we analyzed the performance of RBP-based TL repression in two I1-FFL circuit designs, TL-TX-TL and TX-TL-TL, to explore the effect of regulatory mechanisms at the TL level on pulse generation and circuit dynamics.

For the TL-TX-TL circuit, transcription of RBP at node Y is driven by a weak J23116 promoter, where the translation for RBP is activated by THS trigger from node X. The transcription of GFP output at node Z is activated by STAR and expressed under the pLlacO promoter (Figure 3A). To modulate RBP concentrations, three degradation tags (ASV, AAV, LVA) with varying degradation strength were tested. Experimental results demonstrated that a stronger degradation tag (e.g., LVA) would lead to higher GFP expression levels, as compared to weaker degradation tags (e.g., ASV and AAV) (Figure 3B). The same trend was observed in simulation results (Figure 3C). Due to the strong repression by RBP, the circuits with ASV and AAV-tagged RBPs showed little difference in the GFP output trajectory in experiments, whereas small but noticeable differences were observed for simulation results.

**Figure 3.**
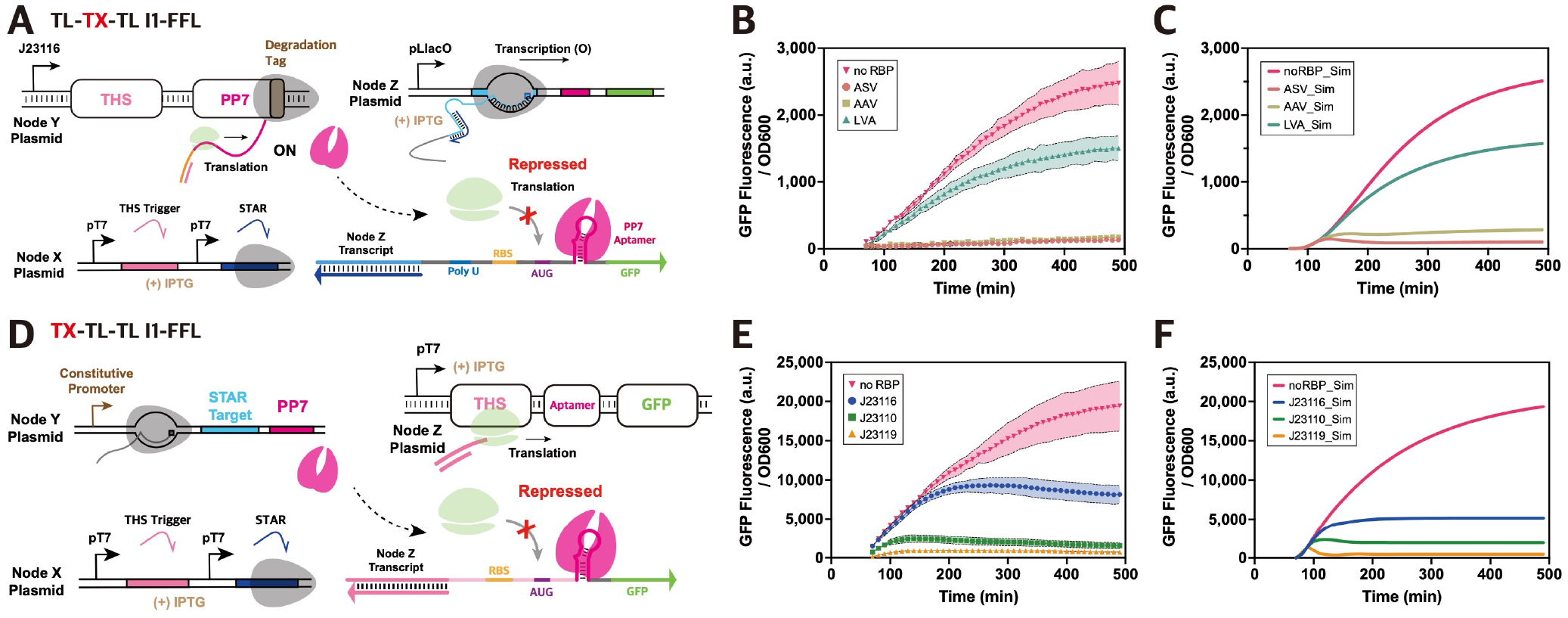
Characterization of I1-FFL Circuits Using RBP as Node Y. (A) Design of TL-TX-TL I1-FFL circuit: Node Y is expressed by THS, and node Z by STAR. RBP (PP7) from node Y binds to the PP7 aptamer in node Z’s transcript, blocking ribosome access and repressing node Z translation. (B) Experimental results for TL-TX-TL circuit for different degradation tags on RBP. GFP fluorescence normalized by OD600 values are plotted, where dots represent the average with shades representing standard deviation of biological triplicate measurements. (C) Simulation results for TL-TX-TL circuit based on the mathematical model. (D) Design of TX-TL-TL I1-FFL circuit: Node Y is expressed by STAR and represses node Z, expressed by THS. (E) Experimental results for TX-TL-TL circuit for different promoters for Y: GFP fluorescence normalized by OD600 values are plotted, where dots represent the average with shades representing standard deviation of biological triplicate measurements. (F) Simulation results for TX-TL-TL cir<u>cuit from the mathematical model.</u>

This suggests insufficient timescale separation between the repression and the activation, where RBP-based repression could prevent accumulation of GFP signal early on, thereby repressing the pulse generation. A pulse-like behavior was observed in simulations with ASV-tagged RBP albeit with quite small amplitude (Figure 3C).

For the TX-TL-TL circuit, transcription of RBP at node Y is activated by STAR and constitutively expressed using different constitutive promoters (Figure 3D). The transcription of GFP output at node Z, driven by the T7 promoter, is repressed by RBP binding to the aptamer within the transcript. Analogous to the TL-TX-TL circuit, no clear pulse was obtained experimentally or in simulations (Figure 3E,F), for the promoter variants of node Y. These findings further support the conclusion that a lack of sufficient timescale separation between the activation and the repression pathways inhibits pulse generation in RBP-based circuits.

To better understand the underlying mechanisms for the lack of pulsed dynamics, we sought to investigate whether it stems from inherent properties of RBP-mediated inhibition or from specific regulatory configurations. This prompted a deeper exploration into the factors influencing circuit dynamics, aiming to uncover potential mechanisms that could enhance pulse generation.

Sensitivity Analysis of TetR and RBP Production and Degradation Rates across Four Circuits. Building on the observed differences in dynamics between TetR and RBP-based circuits, we investigated how key kinetic parameters, such as production and degradation rates, influence circuit behavior. To achieve this, we conducted model-based local sensitivity analysis across the four RNAprotein hybrid I1-FFL circuits (Figures S8-S11). This analysis systematically evaluated the impact of small changes in individual parameters, including transcription rates (*α*), degradation rates (*δ*), binding coefficients (*ω, β, γ*), and decoupling coefficients (*ν*). Given that the primary objective of this work is to investigate how TX and TL regulations shape the I1-FFL circuit dynamics, we focused specifically on the production and degradation rates of TetR and RBP (*α*_*TetR*_, *δ*_*TetR*_, *α*_*RBP*_, *δ*_*RBP*_). These parameters were selected as TetR mediates TX control, while RBP governs TL inhibition, representing the central regulatory mechanisms driving circuit performance.

In the TX-TL-TX circuit, higher TetR production rate *α*_*TetR*_ led to earlier peaks and more prominent pulses in GFP expression (Figure 4A). This behavior likely stems from faster TetR accumulation enabling earlier repression while maintaining a sufficient timescale separation after the activation. Similar effects were observed by decreasing the degradation rate *δ*_*TetR*_, as a lower *δ*_*TetR*_ prolonged TetR presence in the system, reinforcing its inhibitory effect (Figure S12A,S13A). In contrast, in the TL-TX-TX circuit, *δ*_*TetR*_ had a more pronounced impact than *α*_*TetR*_ on pulse formation (Figure 4C). Lower *δ*_*TetR*_ extended the stability of TetR, enabling sustained repression and stronger pulse characteristics, while manipulating *α*_*TetR*_ showed limited influence (Figure S12B,S13B).

**Figure 4.**
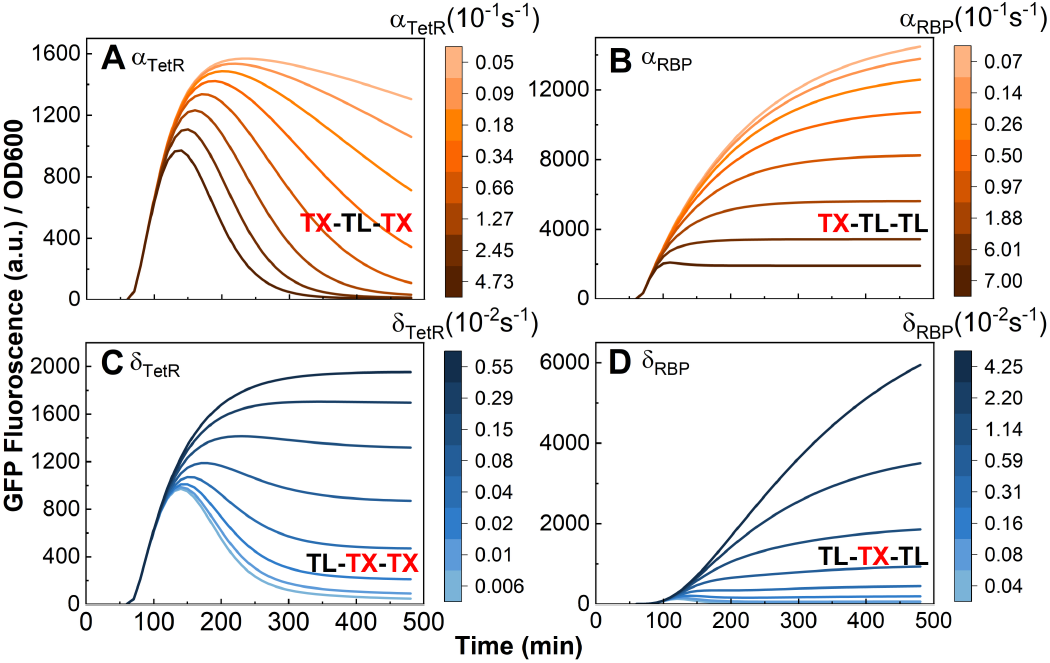
Local Sensitivity Analysis of TetR and RBP Production (α) and Degradation (δ) Rate Across Four I1-FFL Circuit Variants. Parameter values were systematically varied around their fitted nominal values (obtained from previous simulation results) by one order of magnitude above and below. Each parameter range was divided into 8 points using logarithmic spacing to better represent variations across orders of magnitude. The darker the color, the greater the parameter value. (A) TX-TL-TX circuit: *α*_*TetR*_ variation. Higher values lead to earlier, more pronounced GFP peaks. (B) TX-TL-TL circuit: *α*_*RBP*_ variation. Increasing *α*_*RBP*_ improves repression but does not induce clear pulse dynamics. (C) TL-TX-TX circuit: *δ*_*TetR*_ variation. Lower values extend repression, producing distinct pulse dynamics. (D) TL-TX-TL circuit: *δ*_*RBP*_ variation. Reduced *δ*_*RBP*_ increases repression but does not induce clear pulse dynamics.

For RBP-based circuits, simulation results indicated that altering the production rate of RBP (*α*_*RBP*_) and degradation rate (*δ*_*RBP*_) had significant impact on the GFP expression levels but failed to produce discernible pulses (Figure 4B,D). Variations in *δ*_*RBP*_ had a stronger influence on repression strength than changes in *α*_*RBP*_, as longer RBP stability enhanced its inhibitory effects. These findings highlight the limitations of TL regulation in achieving adequate timescale separation for pulse generation, as observed in the TetR circuits (Figures S12C,D,S13C,D).

The observed differences stem from the fundamental differences in repression mechanism and target dynamics between TetR and RBP-based circuits. TetR directly binds to the promoter regions on DNA, effectively halting transcription and ensuring a well-defined timescale separation for pulse generation. In contrast, RBP binds to RNA aptamers on the transcribed RNA, targeting a much larger pool of transcripts compared to the single promoter target of TetR. This disparity in target numbers inherently challenges the efficiency of RBP-based repression, as the abundance of RNA transcripts dilutes the repressor’s impact, requiring more precise tuning to achieve comparable levels of inhibition.

Global sensitivity analysis was conducted to further elucidate these differences, using simultaneous random perturbations across 10,000 parameter combinations and focusing on rise times, pulse widths, heights, and steady states (Figure S14). TetR-based circuits exhibited the highest pulse success rate, while RBP-based circuits showed delayed responses and constrained pulse widths. These findings emphasize the importance of achieving adequate timescale separation for effective pulse generation and highlight the need for strategies to introduce delays in RBP-mediated repression.

Model-Based Strategies for Enhancing Pulse Generation in RBP-Based Circuits. Building on the need for adequate timescale separation identified in our sensitivity analysis, we explored whether introducing delays in RBP-mediated repression could enable robust pulse generation in RBP-based circuits. Computational modeling revealed that artificial delays in RBP production successfully enabled pulse generation in both TL-TX-TL and TX-TL-TL circuits, with TX-TL-TL exhibiting sharper and earlier peaks at shorter delays (80, 120 min), while TL-TX-TL required longer delays (120, 240 min) to produce discernible pulses (Figure S15). These findings underscore the critical role of timing in balancing activation and repression dynamics in RBP-based circuits. Encouraged by these results, we propose translating these model-based insights into experimental strategies, such as using aptamer-based mechanisms to transiently sequester RBP^46^, offering a promising pathway to introduce delays in repression initiation and addressing the limitations observed in RBP-based circuits.

Enhanced Pulse Generation in RBP-Based I1-FFL Circuits Using Aptamer Sponge. To experimentally validate the potential for introducing delays in RBP-mediated repression, we implemented an aptamer-based strategy inspired by the results of our modeling. Previous simulations demonstrated that artificial delays in RBP production enabled robust pulse formation in RBP-based circuits. Building on this, we hypothesized that transiently sequestering RBP could achieve similar temporal effects, delaying repression and promoting pulse generation in experiments.

To test this hypothesis, we designed a single-stranded RNA molecule containing four tandem PP7 aptamer sequences to function as a sponge. This construct, which we refer to as the aptamer sponge, temporarily sequesters RBP, reducing its availability for early-stage binding to transcripts from node Z and mimicking the effect of a delay. We co-transformed this aptamer sponge expressing plasmid with TL-TX-TL and TX-TL-TL I1-FFL circuits and conducted a time course analysis (Figures 5,S16). For the TL-TX-TL circuit, overall GFP expression was low, and distinct pulse-like behavior was not observed, indicating the need for further tuning of parameters. Fortunately, for the TX-TL-TL circuit, we detected a pulse generation for appropriate levels of aptamer sponge expression at 0.2% arabinose in combination with relatively strong promoters for node Y. For instance, when node Y was driven by J23110 promoter, the aptamer sponge expression resulted in a clear pulse, with GFP levels peaking around 150 min and subsequent decay, marking a noticeable pulse production unlike the circuits without the aptamer sponge.

**Figure 5.**
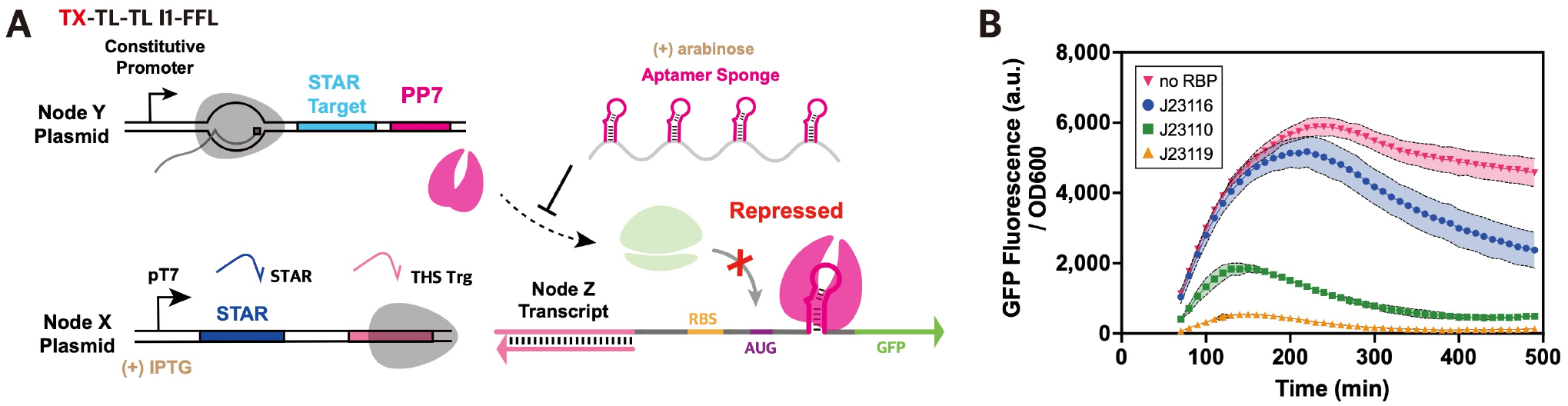
Enhanced Pulse Generation in TX-TL-TL I1-FFL Circuit Using an Aptamer Sponge. (A) Schematic of TX-TL-TL Circuit with Aptamer Sponge: Node Y, regulated by STAR, expresses PP7, which represses node Z (expressed by THS). To create a temporal delay in PP7-mediated repression, an aptamer sponge under the pBAD promoter sequesters PP7 during early stages. (B) Experimental Results of TX-TL-TL Circuit with Aptamer Sponge: GFP expression from node Z is monitored over 490 minutes. Induction with 0.2% arabinose activates the pBAD promoter. GFP fluorescence normalized by OD600 values are plotted, where dots represent the average with shades representing standard deviation of biological triplicate measurements.

The overall GFP expression levels were a bit lower in the presence of aptamer sponge, possibly due to increased cellular burden. Still, early-stage activation remained largely unaffected, suggesting that the aptamer sponge effectively sequestered available RBP early on, mimicking a delay-like effect. This approach provided a temporal separation similar to the effect achieved with aTc in TetR-based circuits, facilitating pulse dynamics in RBP-regulated systems.

Encouraged by the results, we sought to apply the same aptamer sponge strategy to a TL-TL-TL (TL-only) circuit, previously explored only through modeling^37^. In this design, the input node X expressed the THS trigger, with nodes Y and Z configured as in the TLTX-TL and TX-TL-TL circuits, respectively (Figure S17). For appropriate levels of aptamer sponge expression in combination with weaker degradation tags, we observed small pulse peaks for the GFP output, suggesting that obtaining pulsed dynamics in TL-only circuit is feasible but further exploration in parameter space would be required.

Our findings demonstrate that, similar to TetR-based circuits, RBP-based I1-FFL circuits can be optimized to exhibit pulse-like activity through the addition of aptamer sponges. This approach opens up avenues for further development of pulse-generating TL circuits and highlights the potential of aptamer-based delay mechanisms for enhancing the dynamic versatility of I1-FFL circuits in biosensor applications.

## DISCUSSION

In this study, we designed and analyzed four RNA-protein hybrid I1-FFL circuits in *E. coli*, successfully achieving tunable pulse-like dynamics through both TX and TL regulatory mechanisms. By employing RNA-based regulators such as STAR and THS, alongside TetR and RBP as protein-based regulators, we demonstrated the versatility of RNA-protein circuits, capitalizing on RNA’s programmability, rapid signal processing, and the large orthogonal library to lay the foundation for constructing complex biosensing systems.

Our work highlights the novel contributions of RNA-protein hybrid circuits, particularly by incorporating the RBP to achieve TL repression pathways. While TX regulators like TetR are largely confined to target proximal domains to promoters, RBP-based TL inhibition can be flexibly configured with aptamer arrays, offering the potential for enhanced repression efficiency by increasing RBP clustering on target RNAs. Furthermore, aptamer-based regulation supports modular configurations that can be expanded with multiple aptamer-RBP pairs^33^, creating possibilities for complex logic configurations, to reach potential design space beyond simple TX regulators.

Our findings also revealed several distinctions between the TetR-based and the RBP-based repression. TetR demonstrated more efficient repression, likely due to its direct binding to the TetO region on plasmid DNA, effectively blocking transcription^47^. Even with the use of aTc to modulate TetR activity, strong repression was maintained, suggesting robust TX control. In contrast, while RBP-based regulation showed a significant repression, indicating its potential for tight TL control, the experiments showed a relatively weaker repression as compared to TetR. Presumably, RBP that needs to target the aptamers within the transcribed RNAs, which are much more abundant than the corresponding DNA templates, face more challenges to achieve tight control on all of its targets. To potentially increase the repression efficiency of RBP, strategies such as expanding aptamer arrays or combining different aptamers on a single transcript could enhance the clustering and functional efficacy of RBPs.

Despite the dynamic circuit examples demonstrated here, current limitations include the relatively small number of well-characterized RBP-aptamer pairs^48^, which restricts the combinatorial design space of these circuits. Future work could leverage SELEX (Systematic Evolution of Ligands by EXponential enrichment)^49^ to broaden the library of aptamers, thereby expanding the utility of RBP-based repression and creating a more diverse toolkit for TL control. Similarly, both STAR and THS regulators pose their own challenges, such as potential leakage in THS and comparatively low ON-state expression levels for STAR^50^, which can impact circuit performance. These issues could be mitigated by more extensive library characterization and strategic modeling to optimize signal integrity.

This work also demonstrated that circuit dynamics can be finely tuned by adjusting promoter strengths and degradation rates. For instance, exploring a wider range of synthetic promoters, and testing additional synthetic degradation tags^51, 52^can improve circuit robustness in more complex biosensing environment. Furthermore, incorporating elements from the ROSALIND biosensor system^53^, which detects small molecules using transcription factors in cell-free environment, could further improve the applicability of these circuits for specific biosensing tasks. For example, proteins from the broader TetR family^54^ or light-sensitive proteins like EL222^55^ could be incorporated to introduce new stimuli, such as light, for more versatile biosensing applications. Similarly, expanding the RBPs beyond PP7, such as MS2 or Qβ^33^, could offer additional options for regulating gene expression at the TL level, enhancing the modularity of the circuit designs.

Furthermore, the flexibility inherent in RNA-protein hybrid circuits suggests exciting avenues for more sophisticated applications. These circuits lay the groundwork for developing more complex cellular systems where multiple I1-FFL circuits could operate simultaneously^56^, capturing signals across diverse time scales and responding to dynamic environmental inputs. By integrating these distinct timescale responses, a single system could achieve rapid and delayed responses, enhancing adaptability for real-time sensing or environmental monitoring. Future studies could explore multi-pathway circuits that adapt to specific signals or conditions, such as clustered RBP condensates^57^, thereby extending the utility of these systems to stress response and metabolic engineering applications.

Ultimately, our findings underscore the promise of RNA-protein hybrid circuits as flexible, adaptable systems for synthetic biology. By expanding RNA-based regulation and leveraging the unique properties of multi-level I1-FFL circuits, this work lays a foundation for future biosensing systems capable of robust and responsive behavior across a variety of applications, including environmental sensing, complex decision-making, and metabolic pathway optimization.

## METHODS

### Plasmid Construction and *E. coli* Strains Used

Plasmids were constructed using backbones from commercial vectors pET15b, pCDFDuet, pCOLADuet, and pACYCDuet (EMD Millipore, Billerica, MA, USA). The input node X used a pET15b plasmid backbone, while the intermediate node Y and the output node Z used pCDFDuet and pCOLADuet, respectively. The aptamer sponge was constructed in a pACYCDuet plasmid backbone. Constructs were assembled using PCR, Gibson assembly^38^, and round-the-horn site-directed mutagenesis^39^. DNA oligonucleotides were synthesized by Bionics. The RNA regulatory sequences were taken from the library of STAR^12^, THS^13^, and PP7 aptamers^40^. Final constructs were cloned in *E. coli* DH5α for validation via DNA sequencing. Detailed protocols and parts sequences are provided in Supplementary Information.

### Cell Culture and Microplate Reader Time Course Analysis

All *E. coli* cultures were grown on LB agar plates with appropriate antibiotics for selection, then single colonies were inoculated in liquid LB with shaking. Overnight cultures were diluted and grown to log phase before induction. Inducers included 0.1 mM β-d-1-thiogalac-topyranoside (IPTG) for circuits with TetR and RBP-based repression. For TetR-based circuits, anhydrotetracycline (aTc) was also used at different concentrations (100, 50, 20, and 0 ng/mL), while circuits utilizing RBP were treated with arabinose at different concentrations (0.2%, 0.1%, 0.05%, 0.025%) to assess pulse dynamics. GFP fluorescence and optical density at 600 nm (OD600) were measured at 10-minute intervals using a Synergy H1 microplate reader (BioTek, Winooski, VT, USA) over an 8-hour incubation period. Further details on the culture conditions and inducer concentrations can be found in Supplementary Information.

### Microplate Reader Analysis for Spacer Optimization

For optimization of spacer sequences in RBP-regulated circuits, plasmids expressing PP7 or control plasmids lacking PP7 were co-transformed in *E. coli* BL21(DE3) cells with plasmids containing aptamer-based spacer variants in the output node Z. Cells were grown to log phase, induced with 0.1 mM IPTG, and fluorescence was measured after 3.5 hours. The complete set of experimental results on spacer optimization is provided in Supplementary Information.

Mathematical Modeling. The ODE models for the circuits were developed in MATLAB. The models were parameterized and fitted using experimental GFP fluorescence output data with MATLAB *fminsearch* and *fmincon* functions, with sensitivity analyses performed on key kinetic parameters. All MATLAB scripts and protocols for model construction are available on GitHub (https://github.com/SimraShoaib/I1-FFLcircuits). A complete description of the modeling methodology including detailed sensitivity analysis is provided in Supplementary Information.

## ASSOCIATED CONTENT

### Supporting Information

The Supporting Information is available free of charge on the ACS Publications website.

Experimental procedures, supplementary tables, ODE-based modeling and supplementary figures (PDF)

## AUTHOR INFORMATION

### Authors

Seongho Hong - Department of Life Sciences, Pohang University of Science and Technology, Pohang 37673, Republic of Korea;

Syeda Simra Shoaib - Cain Department of Chemical Engineering, Louisiana State University, Baton Rouge, Louisiana 70803, United States;

### Author Contributions

S.H. and S.S.S. contributed equally as co-first authors. S.H. conducted experiments and contributed to writing and discussion. S.S.S. led the modeling work and played a key role in writing the manuscript. M.F., X.T., and J.K. were involved in research planning, discussion, and manuscript writing, review, and editing. All authors reviewed and approved the manuscript.

### Notes

The authors declare no competing financial interest.

## ACKNOWLEDGMENT

This research was supported by Korea Basic Science Institute (National Research Facilities and Equipment Center) grant funded by the Ministry of Education (2021R1A6C101A390); FoodTech RnD Center Development and Support Program through the GBTP (Gyeongbuk Technopark) funded by Gyeongsangbukdo and Pohang city (GBTP2023129001); Korea Health Technology R&D Project funded by the Korea Health Industry Development Institute (KHIDI) (RS-2023-00304637); a synthetic biology grant funded by Gyeongsangbukdo and Pohang city; High Value-added Food Technology Development Program through the Korea Institute of Planning and Evaluation for Technology in Food, Agriculture and Forestry (IPET) funded by the Ministry of Agriculture, Food and Rural Affairs (MAFRA) (RS-2024-00403998); a grant of the Korea Institute of Planning and Evaluation for Technology in Food, Agriculture and Forestry (IPET) through Agriculture and Food Convergence Technologies Program for Research Manpower development Program, funded by Ministry of Agriculture, Food and Rural Affairs (MAFRA) (RS-2024-00402136) (to J.K.). S.S. and X.T. were supported by the startup fund from the Cain Department of Chemical Engineering at Louisiana State University, and the US NSF grant #2223720.

