## Supplementary Information for "Multi-Level Regulation in RNA-Protein Hybrid Incoherent Feedforward Loop Circuits for Tunable Pulse Dynamics in *Escherichia coli*"

§These authors contributed equally.

### Table of Contents

#### 1. Experimental Procedures


1.1 Plasmid Construction and *E.coli* Strains Used p 3
1.2 Cell Culture and Microplate Reader Time Course Analysis p 3
1.3 Microplate Reader Analysis for Spacer Optimization p 3
1.4 Mathematical Modeling p 4


#### 2. Supplementary Tables


Table S1. Plasmids Used for the Experiment and Their Associated Sequences p 5
Table S2. Experimental Results for PP7 Aptamer Spacer Optimization in STAR p 6
Table S3. Experimental Results for PP7 Aptamer Spacer Optimization in THS p 7
Table S4. TX-TL-TX I1-FFL Parameters across Different aTc Conditions p 7
Table S5. TL-TX-TX I1-FFL Parameters across Different aTc Conditions p 8
Table S6. TL-TX-TL I1-FFL Parameters p 9
Table S7. TX-TL-TL I1-FFL Parameters p 10


#### 3. ODE-Based Modeling


3.1 ODE for TX-TL-TX I1-FFL p 11
3.2 ODE for TL-TX-TX I1-FFL p 12
3.3 ODE for TL-TX-TL I1-FFL p 12
3.4 ODE for TX-TL-TL I1-FFL p 13


#### 4. Supplementary Figures


Figure S1. Plasmid Maps Used in I1-FFL Circuits p 14
Figure S2. Comparison of Promoter ON/OFF Ratios in STAR p 15
Figure S3. Optimization of IPTG Concentration in I1-FFL Circuits Using RBP as Node Y p 15
Figure S4. Comparison of Model Predicted TetR across Different Promoter/Tag Variant in TetR-based Circuits p 16
Figure S5. Experimental Results of TetR-Based I1-FFL Circuits Under Varying aTc Concentrations p 16
Figure S6. Modeling Results of TetR-Based I1-FFL Circuits Under Varying aTc Concentrations p 17
Figure S7. Design and Optimization of PP7 Aptamer-Integrated Riboregulator p 17
Figure S8. Local sensitivity analysis of Circuit TX-TL-TX parameters p 18
Figure S9. Local sensitivity analysis of Circuit TL-TX-TX parameters p 19
Figure S10. Local sensitivity analysis of Circuit TL-TX-TL parameters p 20
Figure S11. Local sensitivity analysis of Circuit TX-TL-TL parameters p 21
Figure S12. Sensitivity Analysis of TetR/RBP Production Rate across Four I1-FFL Circuit Configurations p 22
Figure S13. Sensitivity Analysis of TetR/RBP Degradation Rate across Four I1-FFL Circuit Configurations p 23
Figure S14. Global sensitivity analysis demonstrating the dynamics achievable in each circuit p 24
Figure S15. Pulse generation performance in RBP circuits under simulated RBP production delays p 24
Figure S16. Effect of Aptamer Sponge on Pulse Generation in I1-FFL circuits using RBP as a Node Y p 25
Figure S17. Evaluation of Aptamer Sponge on Pulse Generation in TL-TL-TL(TL-only) I1-FFL Circuit p 26


#### 5. References

### 1. Experimental Procedures

#### 1.1 Plasmid Construction and *E. coli* Strains Used

Backbones for the plasmids were taken from the commercial vectors pET15b, pCDFDuet, pCOLADuet and pACYCDuet (EMD Millipore, Billerica, MA, USA). Node X was constructed in pET15b. Node Y and node Z were constructed in pCDFDuet and pCOLADuet. Aptamer sponge was constructed in pACYCDuet. All constructs were cloned via PCR, Gibson assembly[1] and round-the-horn site-directed mutagenesis[2]. DNA templates for all four circuits were assembled from single-stranded DNAs purchased from Bionics. The RNA regulatory sequences were chosen as follows: the STAR-target pair AD1.A5-AD1.S5[3], the THS-trigger pair ACTS_TypeII_N3[4], and the PP7 aptamer sequence PP7-LSm[5]. Synthetic DNA strands were amplified using PCR to generate double-stranded DNAs. The resulting DNAs were then inserted into plasmid backbones using about 30-bp homology domains via Gibson assembly. The promoter was changed from pT7 to other promoters (pLlacO, J23116, J23110, J23119, and pT7(TetO)), and degradation tags were added using round-the-horn site-directed mutagenesis. All plasmids were cloned in the *E.coli* DH5a strain and validated through DNA Sequencing. GFPmut3b-ASV was used as the reporter. This GFP is GFPmut3b with an ASV degradation tag[6]. TetR and PP7 coat protein (PP7) were used with three different degradation tags (ASV, AAV and LVA). PP7 was introduced into the Y node through Gibson assembly, from pET283xFlagPP7CPHis (Addgene plasmid no. 28174). Plasmids were purified using Enzynomics EZ-Pure Plasmid Prep Kit (Catalog number: EP101-200N). Plasmids were transformed into strains via chemical transformation. *In vivo* tests were conducted using the *E. coli* BL21 DE3 strain for all circuits. Sequences of elements commonly used in the plasmids are provided in Table S1 and Figure S1.

#### 1.2 Cell Culture and Microplate Reader Time Course Analysis

Transformed cells were cultured on Luria-Bertani (LB) agar plates (1.5% agar) and then single colonies were inoculated into 500 μL LB liquid medium supplemented with appropriate antibiotics: pCOLADuet (50 μg/mL Kanamycin), pCDFDuet (100 μg/mL Spectinomycin), pET15b (100 μg/mL Ampicillin) and pACYCDuet (25 μg/mL Chloramphenicol). These cells were grown overnight (~16 h) in 2mL 96-well plates with shaking at 800 r.p.m. and 37 °C. Overnight cultures were diluted 1/100 in 500 μL LB liquid medium supplemented with suitable antibiotics and incubated for 80 minutes. In circuits where TetR serves as the inhibitory protein, treatment with 0.1 mM Isopropyl β-d-1-thiogalactopyranoside (IPTG) and anhydrotetracycline (aTc) was administered. aTc was treated at four different concentrations: 100 ng/mL, 50 ng/mL, 20 ng/mL, 0 ng/mL. In the case of circuits utilizing RBP as an inhibitory protein, only treatment with 0.1 mM IPTG was applied. For the pulse-enhanced I1-FFL circuits, 0.1 mM IPTG and Arabinose were treated. Arabinose was treated at four different concentrations: 0.2%, 0.1%, 0.05%, 0.025%. An aliquot of 200 μL of inducer-treated cells was added per well on a 96-well Black Plate (SPL Life Sciences, Pocheon, Korea). Plates were incubated at 37 °C with double-orbital shaking in a Synergy H1 microplate reader (BioTek, Winooski, VT, USA) running Gen5 3.08 software. GFP Fluorescence (excitation: 479 nm, emission: 520 nm) and OD600 were measured at 10-min intervals during incubation. All circuits were incubated for 8 h (49 cycles). GFP fluorescence levels were normalized as follows: GFP fluorescence for LB blank was subtracted and the resulting value was divided by OD600. The number of replicates was three for all circuits. This experimental setup corresponds to the data presented in Figure 2, Figure 4, and Figure S3, which display the GFP fluorescence and growth dynamics in response to varying levels of IPTG and aTc for TetR and RBP-based circuits.

#### 1.3 Microplate Reader Analysis for Spacer Optimization

Plasmids expressing PP7 (pCDFDuet), or control plasmids lacking PP7, were co-transformed into *E. coli* BL21(DE3) cells alongside plasmids containing node Z spacer variants integrated with aptamers (pCOLADuet). Additionally, plasmids expressing the corresponding trigger RNA (pET15b) were co-transformed to activate each node Z. After plating on LB agar supplemented with Spectinomycin (100 μg/mL), Kanamycin (50 μg/mL), and Ampicillin (100 μg/mL), single colonies were inoculated in LB medium containing the same antibiotics and incubated overnight at 37 °C with shaking. Following a 1/100 dilution of the overnight cultures into fresh LB medium containing Spectinomycin, Kanamycin, and Ampicillin, cells were grown for 80 minutes, and 0.1 mM IPTG was added. Cells were incubated for an additional 3.5 hours before measuring GFP fluorescence and OD600 using a Synergy H1 microplate reader. GFP fluorescence values were divided by OD600 for comparison of the spacer variants. All experiments were performed in three replicates, with detailed results recorded in Table S2 and Table S3. This experiment corresponds to the data presented in Figure 3, which investigates the effect of spacer length on gene expression modulation by PP7.

#### 1.4 Mathematical Modeling

The GFP concentration was simulated by solving mathematical ODE models using the MATLAB 'odes23tb' solver. This involved constructing an ODE model using knowledge of the system and estimating the unknown parameters of the models by minimizing the error between simulated and experimental data. All four model circuits were fitted to the experimental measurements using MATLAB built-in functions. Initially, effective starting points for the parameters were determined using 'fminsearch', and then all parameters were constrained within reasonable bounds using 'fmincon'. For the sensitivity analysis, 16 evenly spaced numbers between the bounds for the respective kinetic parameters were generated. All MATLAB scripts related to this work are available on GitHub ( https://github.com/SimraShoaib/I1-FFLcircuit).

### 2. Supplementary Tables

| Name | Circuit | Variant | Architecture | Link |
|---|---|---|---|---|
| Node X | All | | pT7-**THS Trigger**-T7term-pT7-**STAR**-T7term-(Bla Promoter)-AmpR-pBR322 origin-backbone | https://benchling.com/s/seq-e8TbCJKaxZq6MtlL5G91?m=slm-bC0TYE5Xd1tgo62rMTWj |
| Node Y | TX-TL-TX (C1) | **J23116** STAR TetR | **J23116-STAR Target**-Linker-RBS-**TetR**-ASV-T7term-SpecR-(Bla Promoter)-CloDF13 origin-backbone | https://benchling.com/s/seq-uHrKCx5IF2lzw6D1cv2v?m=slm-f1B9OJd5mPqUvCpW3Wuc |
| | | **J23110** STAR TetR | **J23110-STAR Target**-Linker-RBS-**TetR**-ASV-T7term-SpecR-(Bla Promoter)-CloDF13 origin-backbone | https://benchling.com/s/seq-2z6qHKPV3NtggAevEBLl?m=slm-hJ3jmCU3lEsXDpoVPJnk |
| | | **J23119** STAR TetR | **J23119-STAR Target**-Linker-RBS-**TetR**-ASV-T7term-SpecR-(Bla Promoter)-CloDF13 origin-backbone | https://benchling.com/s/seq-EQDhmga7CKbDS2QCzBem?m=slm-XK6H2buH5nAJEe1Ezh6a |
| | TL-TX-TX (C2) | THS TetR **ASV** | J23116-**THS(RBS)**-Linker-**TetR-ASV**-T7term-SpecR-(Bla Promoter)-CloDF13 origin-backbone | https://benchling.com/s/seq-AT9HAzXmOxEgt4yptCMs?m=slm-ZgQenwWLOFrUn3jF2cH8 |
| | | THS TetR **AAV** | J23116-**THS(RBS)**-Linker-**TetR-AAV**-T7term-SpecR-(Bla Promoter)-CloDF13 origin-backbone | https://benchling.com/s/seq-cI3PeUrxGt4Imgk2kgrc?m=slm-3wa1OelHdbIHfxdbqTxZ |
| | | THS TetR **LVA** | J23116-**THS(RBS)**-Linker-**TetR-LVA**-T7term-SpecR-(Bla Promoter)-CloDF13 origin-backbone | https://benchling.com/s/seq-tE29PFIpqVSyMJjo80z5?m=slm-dBovNeOZyhnvuWxu8y6r |
| | TL-TX-TL (C3) | THS PP7 **ASV** | J23116-**THS(RBS)**-Linker-**PP7-ASV**-T7term-SpecR-(Bla Promoter)-CloDF13 origin-backbone | https://benchling.com/s/seq-zKb5LCeeZfkdSbn4R5tb?m=slm-fH0gZgnMdmbSLReT1brv |
| | | THS PP7 **AAV** | J23116-**THS(RBS)**-Linker-**PP7-AAV**-T7term-SpecR-(Bla Promoter)-CloDF13 origin-backbone | https://benchling.com/s/seq-PJZR9jXvQ4PS1wvIavIM?m=slm-XuGINaZWzaIPu16JpruD |
| | | THS PP7 **LVA** | J23116-**THS(RBS)**-Linker-**PP7-LVA**-T7term-SpecR-(Bla Promoter)-CloDF13 origin-backbone | https://benchling.com/s/seq-TG2GkzogrDr2hwA7noIu?m=slm-51yxLlNH7ZEtD9y283MT |
| | TX-TL-TL (C4) | **J23116** STAR PP7 | **J23116-STAR Target**-Linker-RBS-**PP7**-6xHis-T7term-SpecR-(Bla Promoter)-CloDF13 origin-backbone | https://benchling.com/s/seq-Udi8WWE2H5ianJPJcqry?m=slm-FHY8WGtCI5CIZTjhV8qj |
| | | **J23110** STAR PP7 | **J23110-STAR Target**-Linker-RBS-**PP7**-6xHis -T7term-SpecR-(Bla Promoter)-CloDF13 origin-backbone | https://benchling.com/s/seq-ytPZniDWtN9gbr6pq81F?m=slm-cL6JMi9hUtwytsvhdqxX |
| | | **J23119** STAR PP7 | **J23119-STAR Target**-Linker-RBS-**PP7**-6xHis -T7term-SpecR-(Bla Promoter)-CloDF13 origin-backbone | https://benchling.com/s/seq-OmVpTGYM4Smrxi9OVWTt?m=slm-S0KHQ5sXjovEtR8WtwmB |
| | TetR based No TetR | | pT7-Linker-TetR-ASV-T7term-SpecR-(Bla Promoter)-CloDF13 origin-backbone | https://benchling.com/s/seq-cQAT4jEQkjTtNU2oj9YV?m=slm-aUt5dAzFSsclI2T8oEK6 |
| | PP7 based No PP7 | | J23116-PP7-6xHis-T7term-SpecR-(Bla Promoter)-CloDF13 origin-backbone | https://benchling.com/s/seq-979REh8ct8KnUqY1RBt5?m=slm-MJP9Ac5YnzteTrGKcKKq |
| Node Z | TX-TL-TX (C1) | | **pT7tetO-THS(RBS)**-Linker-**GFPmut3b**-ASV-T7term-KanR-(Bla Promoter)-ColA origin-backbone | https://benchling.com/s/seq-pUoIpunrHSKnEtNkg6uj?m=slm-Pf7995kyUYthPBDAEcQ6 |
| | TL-TX-TX (C2) | | **pT7tetO-STAR Target**-Linker-(RBS)-**GFPmut3b**-ASV-T7term-KanR-(Bla Promoter)-ColA origin-backbone | https://benchling.com/s/seq-Z51Jb6tO3mJSxjvM3EUV?m=slm-jogJDkqVhyO7wSEBILUV |

| | | | | |
|---|---|---|---|---|
| | TL-TX-TL (C3) | | **pLlacO**- **STAR Target**-Linker-(RBS)-**PP7_LSm-GFPmut3b**-ASV-T7term-KanR-(Bla Promoter)-ColA origin-backbone | https://benchling.com/s/seq-fimlZ8OCTs5whBH5kWuu?m=slm-6NYRstaWxuc4svIuQOXJ |
| | TX-TL-TL (C4) | | **pT7**-**THS**(**RBS**-**PP7_LSm**)-Linker-**GFPmut3b**-ASV-T7term-KanR-(Bla Promoter)-ColA origin-backbone | https://benchling.com/s/seq-gKAZB6FMInM5opS4G8Z3?m=slm-6ZMWP9fO2YuvtJFjVxGc |
| PP7 + | Spacer Optimization | | J23116-RBS-PP7-6xHis-T7term-SpecR-(Bla Promoter)-CloDF13 origin-backbone | https://benchling.com/s/seq-uzzmEgWeKrzUKvliG8ds?m=slm-1UPO0gGagN3O6O0AdRFu |
| STAR | Spacer Optimization | | **pLlacO**- **STAR Target**-Linker-(RBS)-**GFPmut3b**-ASV-T7term-KanR-(Bla Promoter)-ColA origin-backbone | https://benchling.com/s/seq-pFhFN6XPJOPxGHCmlVBu?m=slm-xNrhwsUXUHXUEAFa2gYo |
| THS | Spacer Optimization | | **pT7**-**THS**(**RBS**)-Linker-**GFPmut3b**-ASV-T7term-KanR-(Bla Promoter)-ColA origin-backbone | https://benchling.com/s/seq-dYiSmguLblFo8hkKEWJY?m=slm-qhbSGyCH5LA4vfdN7U5q |
| Aptamer Sponge | C3 & C4 | | **pBAD**-**Aptamer Sponge**-T7term- CmR-(cat promoter)-p15A origin-LacI-araC-backbone | https://benchling.com/s/seq-uM6YMLzZbApCjcq2zeEy?m=slm-mMNHEt7sSsw7avXsZSgB |

**Table S1. Plasmids Used for the Experiment and Their Associated Sequences**
Plasmids used in this study. Abbreviations are as follows: T7term = T7 terminator, AmpR = ampicillin resistance gene, SpecR = spectinomycin resistance gene, KanR = kanamycin resistance gene, CmR = Chloramphenicol resistance gene. The associated plasmid sequences can be accessed via the provided Benchling links.

| | **GFP Fluorescence (a.u.)/OD600** | | | | | |
|---|---|---|---|---|---|---|
| | **PP7 +** | | **PP7 -** | | | |
| Spacer | Average | STD | Average | STD | ON level (%) | Repression Efficiency (%) |
| 8 nt | 28749.47 | 1640.12 | 128581.57 | 2064.79 | 37.33 | 77.64 |
| **9 nt** | 29394.19 | 832.52 | 424729.95 | 8023.88 | **123.32** | **93.08** |
| 10 nt | 28279.69 | 3358.11 | 335931.74 | 36628.16 | **97.53** | **91.58** |
| 11 nt | 58040.53 | 3003.31 | 251305.00 | 11291.70 | 72.96 | 76.90 |
| 12 nt | 289861.89 | 23794.46 | 485307.67 | 33531.86 | **140.90** | 40.27 |
| 13 nt | 77211.29 | 45755.51 | 338583.53 | 96037.64 | **98.30** | 77.20 |
| 14 nt | 236121.11 | 13992.04 | 245723.43 | 19818.52 | 71.34 | 3.91 |
| STAR | 346524.55 | 14976.73 | 344423.52 | 26128.31 | 100.00 | -0.61 |

**Table S2. Experimental Results for PP7 Aptamer Spacer Optimization in STAR**
This table summarizes the results of optimizing spacer lengths in the PP7 aptamer for gene expression modulation in *E. coli*. ON level represents normalized gene expression in the absence of PP7, using STAR without an aptamer as the 100% reference. Repression efficiency measures the percentage reduction in expression when PP7 is present. Variants with both ON levels and repression efficiency above 90% are highlighted in bold, while insignificant repression (negative values) is marked in gray. The 9 nt variant, with the highest ON level among effective spacers, was selected as the Z node for the TL-TX-TL circuit. Data are the average of three replicates.

| | GFP Fluorescence (a.u.)/OD600 | | | | | |
|---|---|---|---|---|---|---|
| | PP7 + | | PP7 - | | | |
| Spacer | Average | STD | Average | STD | ON level (%) | Repression Efficiency (%) |
| 8 nt | 45900.19 | 2102.91 | 1983893.28 | 24486.50 | 78.84 | **97.69** |
| 9 nt | 80696.25 | 5097.39 | 872740.92 | 68850.60 | 34.68 | **90.75** |
| **10 nt** | 170048.61 | 5876.36 | 2308226.95 | 90306.15 | **91.73** | **92.63** |
| 11 nt | 226188.25 | 53392.80 | 1108793.88 | 168170.99 | 44.07 | 79.60 |
| 12 nt | 786669.40 | 15608.39 | 793505.50 | 71157.50 | 31.54 | 0.86 |
| 13 nt | 1524548.20 | 436318.85 | 2055758.21 | 132369.83 | 81.70 | 25.84 |
| 14 nt | 1057066.91 | 25301.87 | 665570.91 | 520050.64 | 26.45 | -58.82 |
| THS | 2637935.58 | 46783.15 | 2516202.43 | 312480.99 | 100.00 | -4.84 |

**Table S3. Experimental Results for PP7 Aptamer Spacer Optimization in THS**

This table summarizes the optimization results for spacer lengths in the PP7 aptamer used for gene expression modulation in *E. coli*, targeting THS. The ON level represents normalized gene expression in the absence of PP7, with THS (without an aptamer) serving as the 100% reference. Repression efficiency measures the percentage reduction in expression when PP7 is present. Variants where both ON levels and repression efficiency exceeded 90% are highlighted in bold, while insignificant repression (negative values) is marked in gray. The 10 nt variant, being the only spacer to exceed 90% in both metrics, was selected for the Z node in the TX-TL-TL circuit. Data are the average of three replicates.

| | | aTc (ng/ml) | | | | |
|---|---|---|---|---|---|---|
| **no.** | **Parameters** | 50 | 100 | 20 | 0 | **Units** |
| 1 | α_x | 2.0E-03 | 9.7E-04 | 1.1E-03 | 0.42 | $sec^{-1}$ |
| 2 | α_TetR | 0.04 | 0.03 | 0.05 | 0.50 | $sec^{-1}$ |
| 3 | α_z | 0.23 | 0.16 | 0.18 | 5.0E-04 | $sec^{-1}$ |
| 4 | α_GFP | 0.43 | 0.50 | 0.43 | 0.42 | $sec^{-1}$ |
| 5 | δ_x | 3.1E-03 | 6.1E-03 | 1.0E-02 | 4.2E-03 | $sec^{-1}$ |
| 6 | δ_TetR | 1.0E-02 | 9.1E-03 | 1.0E-02 | 5.0E-05 | $sec^{-1}$ |
| 7 | δ_z | 6.1E-04 | 4.8E-04 | 5.9E-04 | 9.3E-03 | $sec^{-1}$ |
| 8 | δ_THS | 1.0E-02 | 5.5E-03 | 1.0E-02 | 8.7E-03 | $sec^{-1}$ |
| 9 | δ_GFP | 7.2E-03 | 9.0E-03 | 1.0E-02 | 1.0E-02 | $sec^{-1}$ |
| 10 | ω_STAR | 4.9E+03 | 4.9E+03 | 4.9E+03 | 1.2E+02 | $M^{-1}$ $sec^{-1}$ |
| 11 | ν_(y_act) | 1.0E-05 | 1.0E-07 | 1.1E-07 | 1.0E-07 | $sec^{-1}$ |
| 12 | ω_THS | 8.0E+02 | 7.0E+02 | 9.0E+02 | 2.3E+02 | $M^{-1}$ $sec^{-1}$ |
| 13 | ω_TetR | 1.0E+06 | 1.0E+06 | 1.0E+06 | 1.3E+02 | $M^{-1}$ $sec^{-1}$ |
| 14 | β_(aTc_TetR) | 1.5E+04 | 1.5E+04 | 1.5E+04 | 1.0E+02 | $M^{-1}$ $sec^{-1}$ |
| 15 | ν_(Pz_rep) | 1.0E-05 | 2.2E-08 | 1.9E-08 | 1.0E-08 | $sec^{-1}$ |
| 16 | ν_(aTc_TetR) | 1.2E-08 | 2.1E-06 | 9.9E-06 | 1.4E-04 | $sec^{-1}$ |
| 17 | Scaling Factor | 2.47 | 2.62 | 2.88 | 3.17 | - |

**Table S4. TX-TL-TX (C1) I1-FFL Parameters across Different aTc Conditions**

| | | aTc (ng/ml) | | | | |
|---|---|---|---|---|---|---|
| **no.** | **Parameters** | 50 | 100 | 20 | 0 | **Units** |
| 1 | α_x | 0.50 | 0.44 | 0.50 | 0.31 | $sec^{-1}$ |
| 2 | α_TetR | 0.19 | 0.29 | 0.50 | 5.2E-03 | $sec^{-1}$ |
| 3 | α_y | 0.36 | 0.25 | 0.39 | 0.50 | $sec^{-1}$ |
| 4 | α_z | 0.50 | 0.46 | 1.5E-03 | 0.15 | $sec^{-1}$ |
| 5 | α_GFP | 0.18 | 0.41 | 0.50 | 0.30 | $sec^{-1}$ |
| 6 | δ_x | 1.7E-03 | 2.3E-03 | 2.6E-03 | 6.4E-03 | $sec^{-1}$ |
| 7 | δ_THS | 2.1E-03 | 3.3E-03 | 1.3E-03 | 4.5E-04 | $sec^{-1}$ |
| 8 | δ_TetR | 1.0E-02 | 5.5E-04 | 9.4E-04 | 5.5E-04 | $sec^{-1}$ |
| 9 | δ_y | 2.1E-03 | 2.6E-03 | 3.2E-03 | 6.0E-03 | $sec^{-1}$ |
| 10 | δ_z | 1.0E-02 | 4.8E-03 | 1.0E-02 | 1.0E-02 | $sec^{-1}$ |
| 11 | δ_y_act | 2.5E-03 | 4.4E-04 | 1.3E-03 | 8.9E-03 | $sec^{-1}$ |
| 12 | δ_GFP | 1.0E-02 | 6.8E-03 | 2.0E-03 | 1.9E-04 | $sec^{-1}$ |
| 13 | ω_STAR | 3.3E+02 | 1.1E+02 | 1.2E+02 | 1.0E+02 | $M^{-1}sec^{-1}$ |
| 14 | ν_(z_act) | 1.0E-05 | 1.8E-04 | 1.0E-04 | 1.3E-04 | $sec^{-1}$ |
| 15 | ω_THS | 5.7E+02 | 1.0E+02 | 2.3E+02 | 1.1E+04 | $M^{-1}$ $sec^{-1}$ |
| 16 | ω_TetR | 1.3E+03 | 1.1E+02 | 1.7E+02 | 1.4E+04 | $M^{-1}$ $sec^{-1}$ |
| 17 | β_(aTc_TetR) | 1.0E+03 | 1.0E+02 | 1.0E+02 | 3.8E+02 | $M^{-1}$ $sec^{-1}$ |
| 18 | ν_(Pz_rep) | 1.0E-05 | 2.0E-04 | 2.0E-04 | 1.3E-08 | $sec^{-1}$ |
| 19 | ν_(aTc_TetR) | 1.0E-05 | 1.3E-04 | 2.0E-04 | 1.0E-08 | $sec^{-1}$ |
| 20 | Scaling Factor | 0.57 | 0.56 | 2.60 | 0.50 | - |

**Table S5. TL-TX-TX (C2) I1-FFL Parameters across Different aTc Conditions**

| no. | Parameters | Value | Units |
|---|---|---|---|
| 1 | $\alpha\_x$ | 0.13 | $sec^{-1}$ |
| 2 | $\alpha\_y$ | 0.50 | $sec^{-1}$ |
| 3 | $\alpha\_RBP$ | 0.50 | $sec^{-1}$ |
| 4 | $\alpha\_z$ | 0.50 | $sec^{-1}$ |
| 5 | $\alpha\_GFP$ | 9.5E-03 | $sec^{-1}$ |
| 6 | $\delta\_x$ | 9.5E-03 | $sec^{-1}$ |
| 7 | $\delta\_THS$ | 4.4E-03 | $sec^{-1}$ |
| 8 | $\delta\_y$ | 1.3E-03 | $sec^{-1}$ |
| 9 | $\delta\_RBP$ | 1.6E-02 | $sec^{-1}$ |
| 10 | $\delta\_z$ | 0.01 | $sec^{-1}$ |
| 11 | $\delta\_GFP$ | 1.7E-03 | $sec^{-1}$ |
| 12 | $\omega\_STAR$ | 1.2E+03 | $M^{-1}$ $sec^{-1}$ |
| 13 | $\nu\_(Pz\_act)$ | 3.0E-04 | $sec^{-1}$ |
| 14 | $\omega\_THS$ | 3.3E+02 | $M^{-1}$ $sec^{-1}$ |
| 15 | $\omega\_RBP$ | 9.2E+02 | $M^{-1}$ $sec^{-1}$ |
| 16 | $\nu\_(Z\_rep)$ | 1.3E-04 | $sec^{-1}$ |
| 17 | $\delta\_(Z\_rep)$ | 9.5E-04 | $sec^{-1}$ |
| 18 | Scaling Factor | 0.01 | - |

**Table S6. TL-TX-TL (C3) I1-FFL Parameters**

| no. | Parameters | Value | Units |
|---|---|---|---|
| 1 | α_x | 0.47 | $sec^{-1}$ |
| 2 | α_RBP | 0.50 | $sec^{-1}$ |
| 3 | α_z | 0.24 | $sec^{-1}$ |
| 4 | α_GFP | 0.28 | $sec^{-1}$ |
| 5 | δ_x | 1.3E-04 | $sec^{-1}$ |
| 6 | δ_THS | 4.9E-03 | $sec^{-1}$ |
| 7 | δ_RBP | 5.1E-03 | $sec^{-1}$ |
| 8 | δ_Z | 5.6E-03 | $sec^{-1}$ |
| 9 | δ_GFP | 6.6E-03 | $sec^{-1}$ |
| 10 | ω_STAR | 1.8E+03 | $M^{-1}$ $sec^{-1}$ |
| 11 | ν_(y_act) | 1.0E-07 | $sec^{-1}$ |
| 12 | ω_THS | 1.0E+02 | $M^{-1}$ $sec^{-1}$ |
| 13 | ω_RBP | 9.56E+03 | $M^{-1}$ $sec^{-1}$ |
| 14 | ν_(z_rep) | 2.7E-08 | $sec^{-1}$ |
| 15 | Scaling Factor | 2.08 | - |
| 16 | δ_(Z_rep) | 1.0E-04 | $sec^{-1}$ |
| 17 | δ_(Z_act) | 1.0E-04 | $sec^{-1}$ |

**Table S7. TX-TL-TL (C4) I1-FFL Parameters**

### 3. ODE-Based Modeling

#### 3.1 ODE for TX-TL-TX (C1) I1-FFL

$$\frac{d[STAR]}{dt} = \alpha_x[P_{xtotal}] - \omega_{STAR}[STAR][P_{yfree}] - \delta_x[STAR] + \nu_{yact}[P_{yactive}]$$

$$\frac{d[THS\ Trg]}{dt} = \alpha_x[P_{xtotal}] - \omega_{THS}[THS\ Trg][Z] - \delta_{THS}[THS\ Trg]$$

$$\frac{d[TetR]}{dt} = \alpha_{TetR}[P_{yactive}] - \omega_{TetR}[TetR][P_{zfree}] - \beta_{aTc_{TetR}}[aTc][TetR] - \delta_{TetR}[TetR] + \nu_{Pz_{rep}}[P_{zrep}] + \nu_{aTc_{TetR}}[aTc{:}\ TetR]$$

$$\frac{d[aTc]}{dt} = -\beta_{aTc_{TetR}}[aTc][TetR] + \nu_{aTc_{TetR}}[aTc{:}\ TetR]$$

$$\frac{d[aTc{:}\ TetR]}{dt} = \beta_{aTc_{TetR}}[aTc][TetR] - \nu_{aTc_{TetR}}[aTc{:}\ TetR]$$

$$\frac{d[P_{zrep}]}{dt} = \omega_{TetR}[TetR][P_{zfree}] - \nu_{Pz_{rep}}[P_{zrep}]$$

$$\frac{d[Z]}{dt} = \alpha_z[P_{zfree}] - \omega_{THS}[THS\ Trg][Z] - \delta_z[Z]$$

$$\frac{d[Z^*]}{dt} = \omega_{THS}[THS\ Trg][Z] - \delta_{Z^*}[Z^*]$$

$$\frac{d[GFP]}{dt} = \alpha_{GFP}[Z^*] - \delta_{GFP}[GFP]$$

$$\frac{d[P_{yactive}]}{dt} = \omega_{STAR}[STAR][P_{yfree}] - \nu_{yact}[P_{yactive}]$$

$$[P_{yfree}] = P_{ytotal} - P_{yactive}$$

$$[P_{zfree}] = P_{ztotal} - P_{zrep}$$

#### 3.2 ODE for TL-TX-TX (C2) I1-FFL

$$\frac{d[STAR]}{dt} = \alpha_x[P_{xtotal}] - \omega_{STAR}[STAR][P_{zfree}] - \delta_x[STAR] + \nu_{z_{act}}[P_{zactive}]$$

$$\frac{d[THS\ Trg]}{dt} = \alpha_x[P_{xtotal}] - \omega_{THS}[THS\ Trg][Y] - \delta_{THS}[THS\ Trg]$$

$$\frac{d[TetR]}{dt} = \alpha_{TetR}[Y^*] - \omega_{TetR}[TetR][P_{zfree} + P_{zactive}] - \beta_{aTc_{TetR}}[aTc][TetR] - \delta_{TetR}[TetR] + \nu_{Pz_{rep}}[P_{zrep}] + \nu_{aTc_{TetR}}[aTc: TetR]$$

$$\frac{d[aTc]}{dt} = -\beta_{aTc_{TetR}}[aTc][TetR] + \nu_{aTc_{TetR}}[aTc: TetR]$$

$$\frac{d[aTc: TetR]}{dt} = \beta_{aTc_{TetR}}[aTc][TetR] - \nu_{aTc_{TetR}}[aTc: TetR]$$

$$\frac{d[Y]}{dt} = \alpha_Y[P_{ytotal}] - \omega_{THS}[THS\ Trg][Y] - \delta_Y[Y]$$

$$\frac{d[Y^*]}{dt} = \omega_{THS}[THS\ Trg][Y] - \delta_{Y^*}[Y^*]$$

$$\frac{d[P_{zrep}]}{dt} = \omega_{TetR}[TetR][P_{zfree} + P_{zactive}] - \nu_{Pz_{rep}}[P_{zrep}]$$

$$\frac{d[P_{zactive}]}{dt} = \omega_{STAR}[STAR][P_{zfree}] - \omega_{TetR}[TetR][P_{zactive}] - \nu_{z_{act}}[P_{zactive}]$$

$$\frac{d[Z]}{dt} = \alpha_z[P_{zactive}] - \delta_z[Z]$$

$$\frac{d[GFP]}{dt} = \alpha_{GFP}[Z] - \delta_{GFP}[GFP]$$

$$[P_{zfree}] = P_{ztotal} - P_{zrep} - P_{zactive}$$

#### 3.3 ODE for TL-TX-TL (C3) I1-FFL

$$\frac{d[STAR]}{dt} = \alpha_x[P_{xtotal}] - \omega_{STAR}[STAR][P_{zfree}] - \delta_x[STAR] + \nu_{z_{act}}[P_{zactive}]$$

$$\frac{d[THS\ Trg]}{dt} = \alpha_x[P_{xtotal}] - \omega_{THS}[THS\ Trg][Y] - \delta_{THS}[THS\ Trg]$$

$$\frac{d[Y]}{dt} = \alpha_Y[P_{ytotal}] - \omega_{THS}[THS\ Trg][Y] - \delta_Y[Y]$$

$$\frac{d[Y^*]}{dt} = \omega_{THS}[THS\ Trg][Y] - \delta_{Y^*}[Y^*]$$

$$\frac{d[RBP]}{dt} = \alpha_{RBP}[Y^*] - \omega_{RBP}[RBP][Z] - \delta_{RBP}[RBP] + \nu_{Z^{rep}}[Z^{rep}] + \delta_z[Z^{rep}]$$

$$\frac{d[P_{zactive}]}{dt} = \omega_{STAR}[STAR][P_{zfree}] - \nu_{z_{act}}[P_{zactive}]$$

$$\frac{d[Z]}{dt} = \alpha_z[P_{zactive}] - \omega_{RBP}[RBP][Z] - \delta_z[Z] + \delta_{RBP}[Z^{rep}] + \nu_{Z^{rep}}[Z^{rep}]$$

$$\frac{d[Z^{rep}]}{dt} = \omega_{RBP}[RBP][Z] - \delta_z[Z^{rep}] - \nu_{Z^{rep}}[Z^{rep}] - \delta_{RBP}[Z^{rep}]$$

$$\frac{d[GFP]}{dt} = \alpha_{GFP}[Z^{free}] - \delta_{GFP}[GFP]$$

$$[P_{zfree}] = P_{ztotal} - P_{zactive}$$

$$Z^{free} = Z - Z^{rep}$$

##### 3.4 ODE for TX-TL-TL (C4) I1-FFL

$$\frac{d[STAR]}{dt} = \alpha_x[P_{xtotal}] - \omega_{STAR}[STAR][P_{yfree}] - \delta_x[STAR] + \nu_{Py_{act}}[P_{yactive}]$$

$$\frac{d[THS\ Trg]}{dt} = \alpha_x[P_{xtotal}] - \omega_{THS}[THS\ Trg][Z] - \delta_{THS}[THS\ Trg]$$

$$\frac{d[P_{yactive}]}{dt} = \omega_{STAR}[STAR][P_{yfree}] - \nu_{Py_{act}}[P_{yactive}]$$

$$\frac{d[RBP]}{dt} = \alpha_{RBP}[P_{yactive}] - \omega_{RBP}[RBP][Z + Z^*] - \delta_{RBP}[RBP] + \nu_{Z_{rep}}[Z_{rep}] + \delta_{Zact}[Z^{rep}] + \delta_Z[Z^{rep}]$$

$$\frac{d[Z]}{dt} = \alpha_z[P_{ztotal}] - \omega_{RBP}[RBP][Z] - \omega_{THS}[THS\ Trg][Z] - \delta_z[Z] + \nu_{z_{rep}}[Z_{rep}] + \delta_{RBP}[Z^{rep}]$$

$$\frac{d[Z^*]}{dt} = \omega_{THS}[THS\ Trg][Z] - \delta_{Zact}[Z^*] - \omega_{RBP}[RBP][Z^*] + \nu_{z_{rep}}[Z_{rep}] + \delta_{RBP}[Z^{rep}]$$

$$\frac{d[Z^{rep}]}{dt} = \omega_{RBP}[RBP][Z + Z^*] - \delta_{Zrep}[Z^{rep}] - \nu_{z_{rep}}[Z_{rep}] - \delta_{RBP}[Z^{rep}] - \delta_{Zact}[Z^{rep}] - \delta_Z[Z^{rep}]$$

$$\frac{d[GFP]}{dt} = \alpha_{GFP}[Z^*] - \delta_{GFP}[GFP]$$

$$[P_{yfree}] = P_{ytotal} - P_{yactive}$$

#### 4. Supplementary Figure

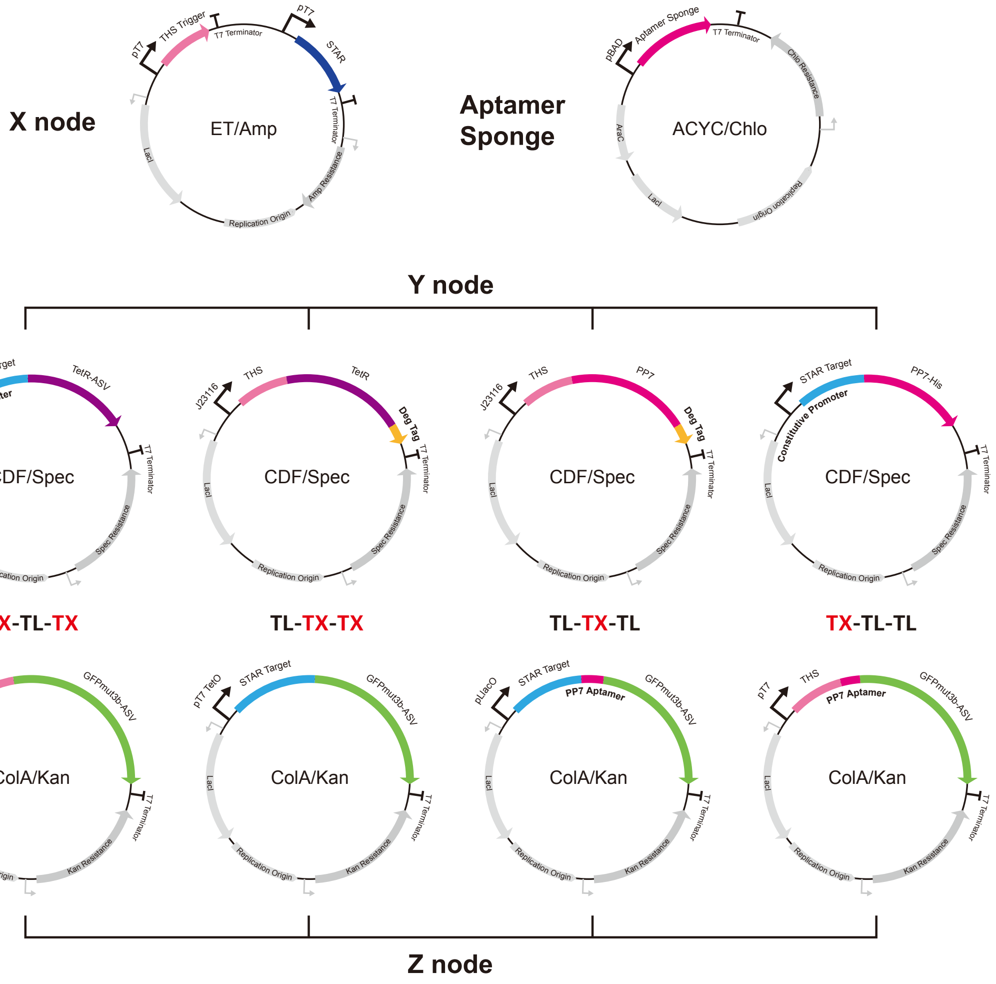


**Figure S1. Plasmid Maps Used in I1-FFL Circuits**

Schematic overview of the plasmid construction used in the I1-FFL synthetic gene circuits. The X node (top) is regulated by the T7 promoter in pET15b, while the Y and Z nodes are regulated by various promoters and aptamer sequences, integrated into pCDFDuet and pCOLADuet plasmids, respectively. Aptamer sponge was constructed in pACYCDuet. The RNA-based regulators STAR, THS, and PP7 aptamers are inserted along with protein-based regulators such as TetR and PP7 coat protein. Promoters and degradation tags were cloned to meet the experimental conditions for optimal regulation of gene expression. Sequences and additional details are provided in Table S1.

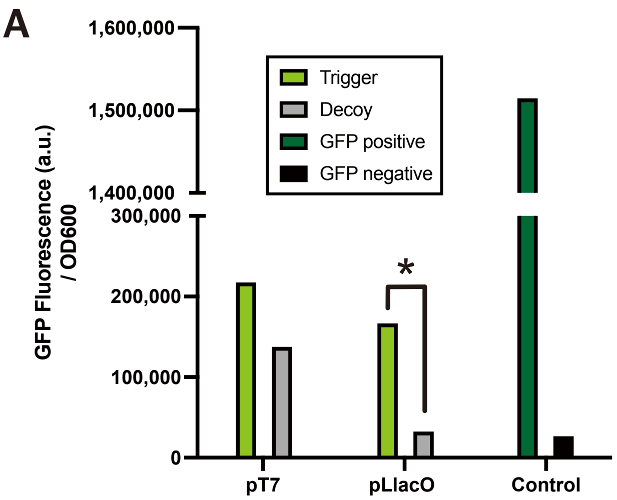


B

| Normalized GFP Fluorescence/OD600 | | |
|---|---|---|
| Promoter | pT7 | pLlacO |
| Trigger | 192034.58 | 199687.30 |
| Decoy | 109702.79 | 5268.49 |
| Fold Change | 1.75 | 26.51 |

**Figure S2. Comparison of Promoter ON/OFF Ratios in STAR**

(A) GFP fluorescence levels (normalized by dividing by OD600) measured under different conditions, both with and without STAR trigger expression, to compare the ON/OFF ratios between the pLlacO and pT7 promoters. Cells containing the pLlacO promoter demonstrate a higher ON/OFF ratio compared to those with the pT7 promoter. Data were collected following the procedures outlined in Experimental Procedures 1.3. Control conditions included BL21 DE3 cells with no plasmid and cells expressing GFP without the STAR Target. Since the experiments were conducted as singlets, no error bars are provided. (B) The normalized GFP fluorescence values for cells under the pLlacO and pT7 promoters, with and without the STAR trigger. Fluorescence values were normalized by subtracting the GFP fluorescence of the negative control and dividing by OD600. The data indicate that the pLlacO promoter achieves a superior ON/OFF ratio, supporting its use in STAR-regulated circuits.

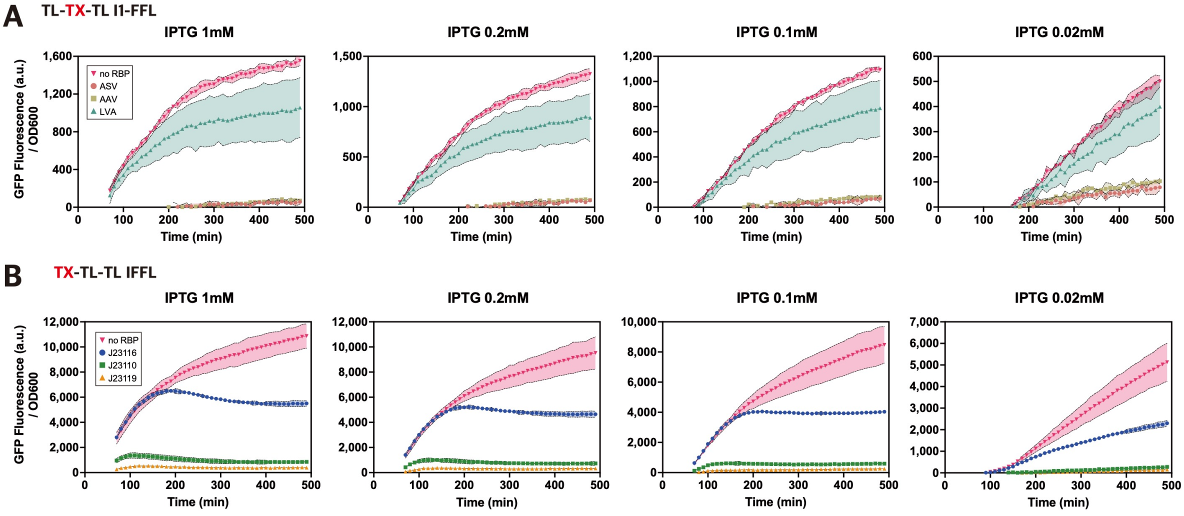


**Figure S3. Optimization of IPTG Concentration in I1-FFL Circuits Using RBP as Node Y**

GFP expression from node Z was monitored across four IPTG concentrations (1 mM, 0.2 mM, 0.1 mM, and 0.02 mM) in both TL-TX-TL and TX-TL-TL I1-FFL circuits to assess circuit response under varying levels of induction. GFP fluorescence and OD600 were measured over 490 minutes. Data are normalized and represent the average of three replicates, with shaded areas indicating standard deviations. (A) TL-TX-TL I1-FFL circuit: PP7 was expressed with different C-terminal degradation tags, each with distinct degradation strengths. Different conditions are indicated by distinct symbols: ASV degradation tag (salmon circles), AAV degradation tag (khaki squares), and LVA degradation tag (teal triangles). Conditions without PP7 are represented by magenta inverted triangles. (B) TX-TL-TL I1-FFL circuit: Node Y was expressed under different promoters: J23116 (blue circles), J23110 (green squares), and J23119 (orange triangles), allowing comparison across promoter strengths. Conditions without PP7 are represented by magenta inverted triangles. For circuits using TetR as node Y, IPTG induction was conducted solely at 0.1 mM, as previously optimized in our prior studies[7], without further concentration comparison.

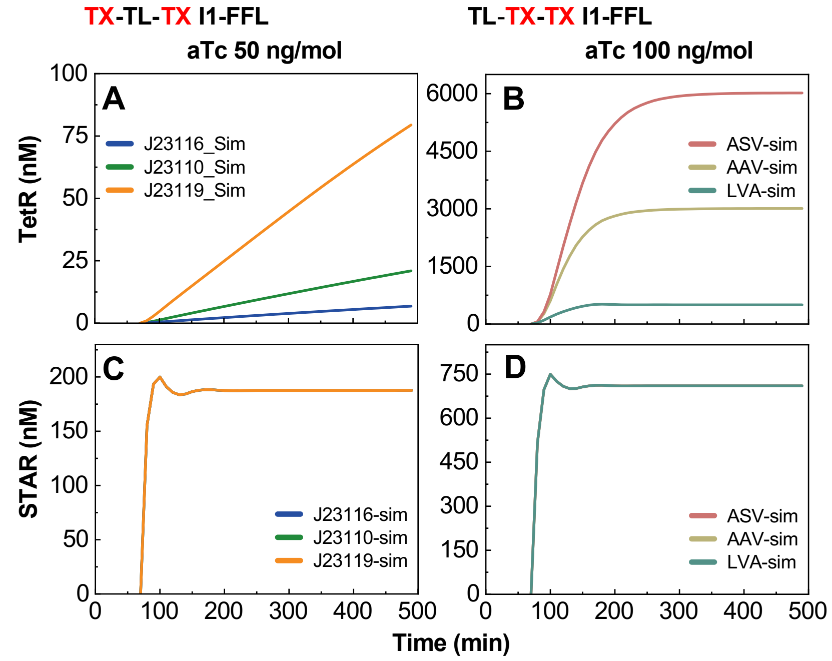


**Figure S4. Comparison of Model Predicted TetR/STAR concentration across Different Promoter/Tag Variant in TetR-based Circuits**

(A) TetR levels decreased more than 10-folds as promoter strength decreased from J23119(strong) to J23116(weakest), indicating a strong inverse relationship between promoter strength and TetR expression. (B) Conversely, TetR concentration increased as the strength of the degradation tag diminished, moving from LVA (strong) to ASV (weakest), highlighting the impact of degradation tag strength on TetR accumulation. (C) STAR concentration remained the same, independent off the promoter strength in the TX-TL-TX circuit. (D) STAR concentration remained constant independent off the protein degradation tag. The concentration difference between TetR and STAR intensifies from a strong degradation tag (LVA) to a weak tag (ASV).

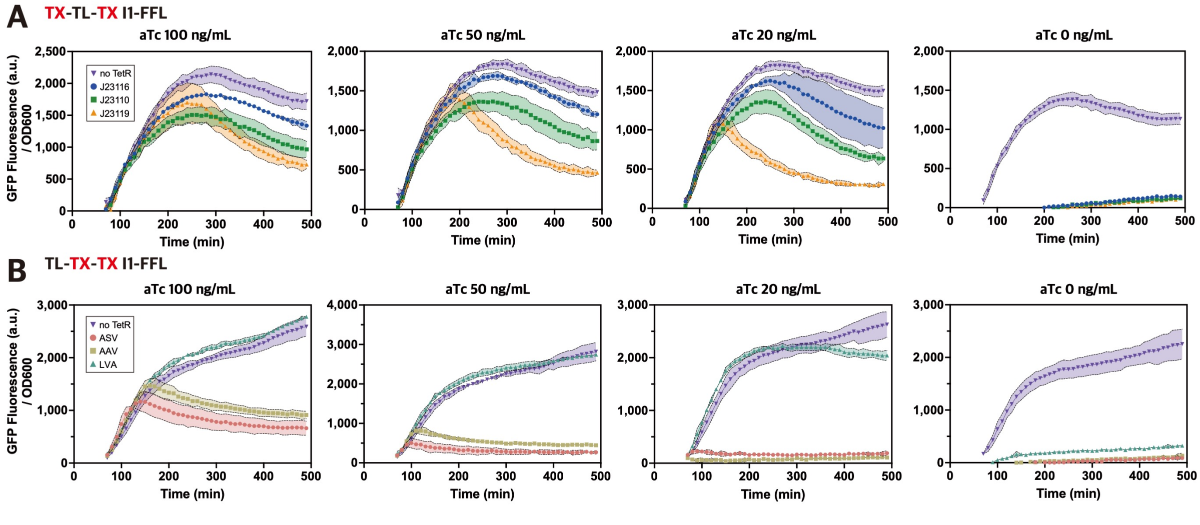


**Figure S5. Experimental Results of TetR-Based I1-FFL Circuits Under Varying aTc Concentrations**

(A) GFP fluorescence response of the TX-TL-TX I1-FFL circuit was measured across four different concentrations of aTc (100 ng/mL, 50 ng/mL, 20 ng/mL, and 0 ng/mL). GFP expression from node Z was monitored for 490 minutes. Data are normalized and represent the average of three replicates, with standard deviations shown as shaded areas. TetR repression was regulated via aTc titration to observe the impact on GFP expression levels. (B)Similar experiments were conducted for the TL-TX-TX I1-FFL circuit, where GFP expression from node Z was tracked under the same aTc concentrations (100 ng/mL, 50 ng/mL, 20 ng/mL, and 0 ng/mL). TetR repression was varied to investigate its effect on translational inhibition. Data are normalized and represent the average of three replicates, with standard deviations depicted as shaded areas.

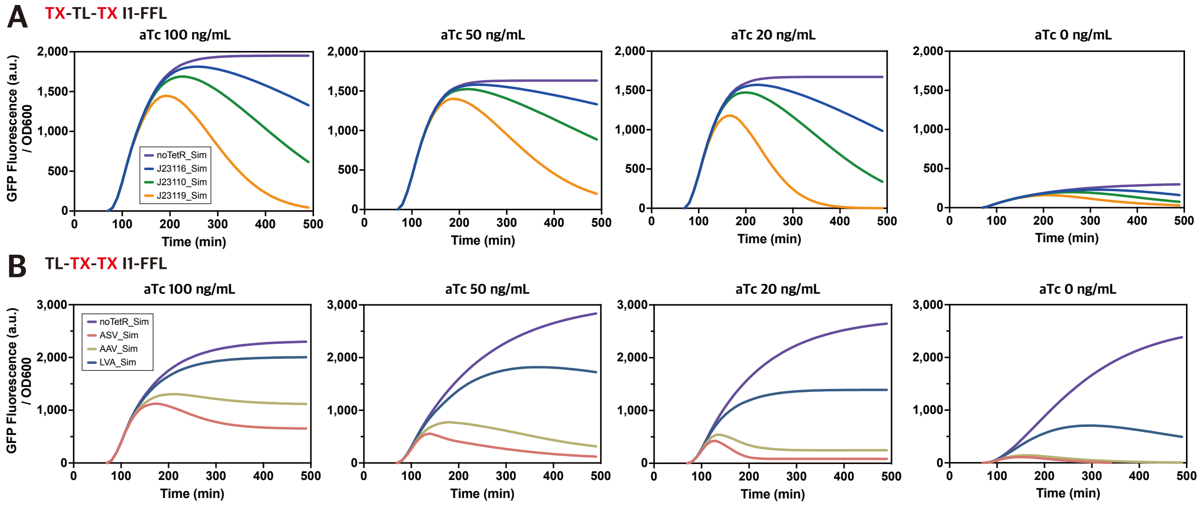


**Figure S6. Modeling Results of TetR-Based I1-FFL Circuits Under Varying aTc Concentrations**
(A) Simulated GFP fluorescence response of the TX-TL-TX I1-FFL circuit across four aTc concentrations (100 ng/mL, 50 ng/mL, 20 ng/mL, and 0 ng/mL) was obtained by minimizing the objective function using the fmincon algorithm in MATLAB. The simulation results capture the dynamic GFP expression from node Z over 490 minutes. (B) Simulated GFP fluorescence response of the TL-TX-TX I1-FFL circuit under identical conditions and optimization approach, showing the impact of aTc-dependent TetR regulation of the TL-TX-TX I1-FFL circuit.

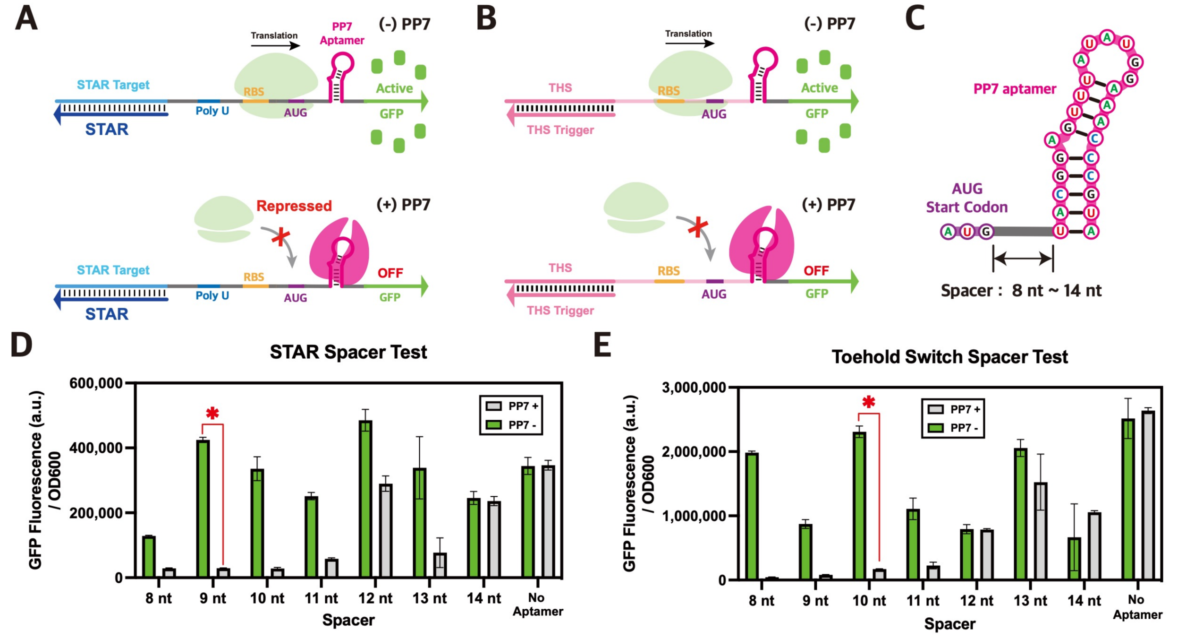


**Figure S7. Design and Optimization of PP7 Aptamer-Integrated Riboregulator**
(A) PP7 aptamer integration with STAR target: In the absence of PP7, STAR activates transcription, enabling normal GFP translation. When PP7 is present, it binds to the PP7 aptamer, blocking ribosome access to the RBS, thus inhibiting GFP expression. (B) PP7 aptamer integration with THS: THS triggers translation of GFP in the absence of PP7. When PP7 is present, it binds to the aptamer, preventing ribosome access and inhibiting GFP expression. (C) Spacer optimization strategy: PP7 aptamer positioned downstream of the start codon with spacers (8-14 nt) to assess repression efficiency. (D, E) Experimental results for spacer optimization with STAR and THS, respectively. Constructs with optimized spacer lengths are marked with a red asterisk. GFP fluorescence normalized by OD600 values are plotted in bar graphs with error bars representing standard deviation of biological triplicate measurements.

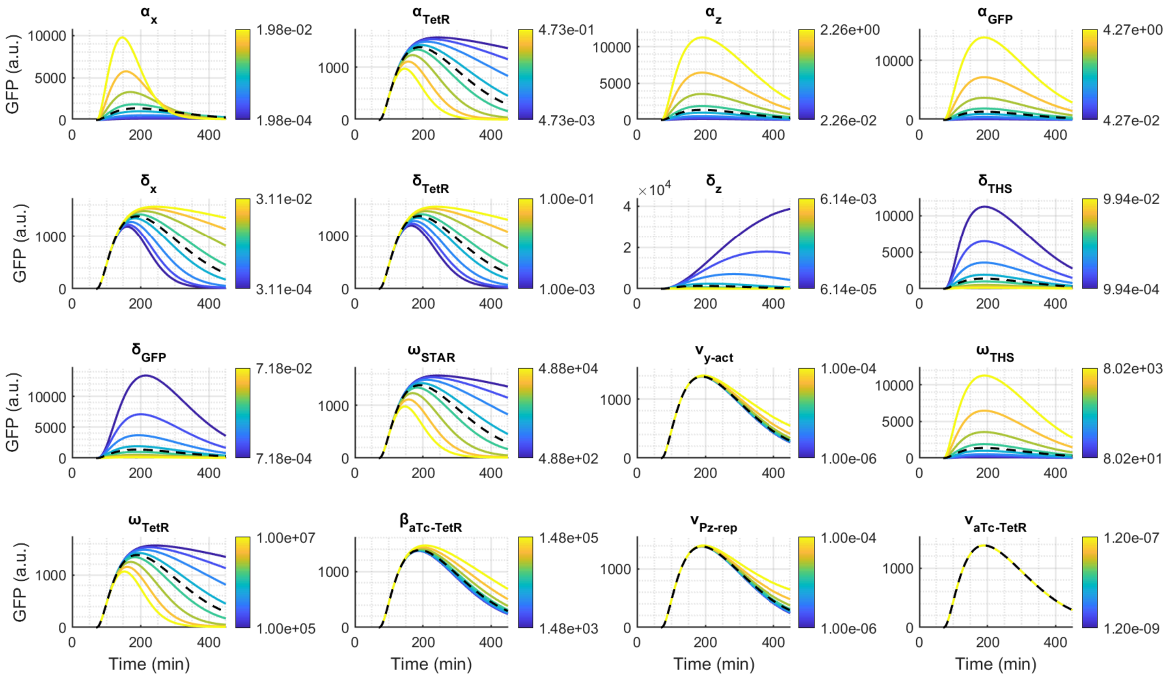

**Figure S8. Local sensitivity analysis of Circuit TX-TL-TX parameters**

Sensitivity analysis of system's output response (GFP expression) to variations in 16 individual parameters while keeping all other parameters at their nominal values. Each subplot represents a different parameter, with colored lines indicating the system response across a range of parameter values (shown in the colorbar, from minimum (blue) to maximum (yellow)).The variation in GFP output (black dashed line) was analyzed by adjusting the underlying parameter values. Parameters analyzed include transcription rates (α), degradation rates (δ), binding coefficients (ω, β, γ), and decoupling coefficients (ν). Time courses demonstrate how perturbations in each parameter affect both the transient dynamics and steady-state behavior of the circuit. Analysis reveals varying degrees of parameter sensitivity, with some parameters (e.g., $\alpha_x$, $\alpha_{TetR}$) showing strong influence on circuit behavior, while others (e.g., $\nu_{Pzrep}$) demonstrate minimal influence.

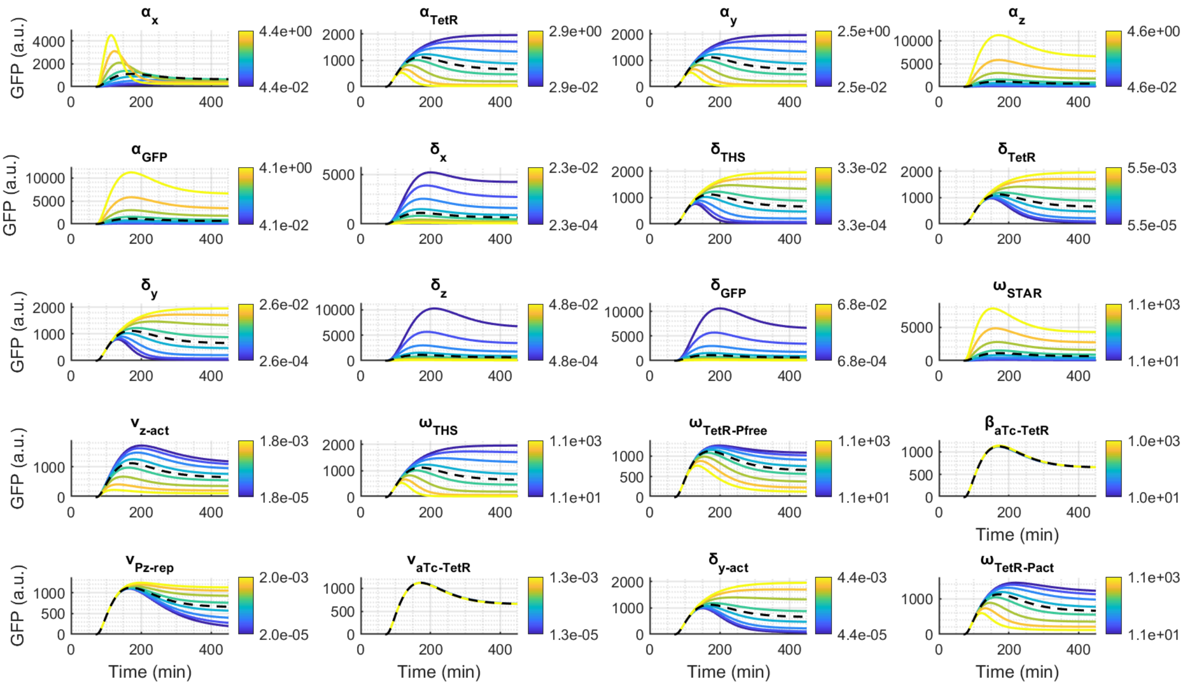


**Figure S9. Local sensitivity analysis of Circuit TL-TX-TX parameters**

Sensitivity analysis of GFP expression in the TL-TX-TX circuit under 20 individual parameter's isolated variations. The variation in GFP output (black dashed line) was analyzed by adjusting the underlying parameter values. Colored lines indicate system responses to individual parameter changes (colorbar: minimum (blue) to maximum (yellow)), with the black dashed line representing the nominal response. Key parameters influencing GFP dynamics include $\delta_x$, $\alpha_{TetR}$ , etc.

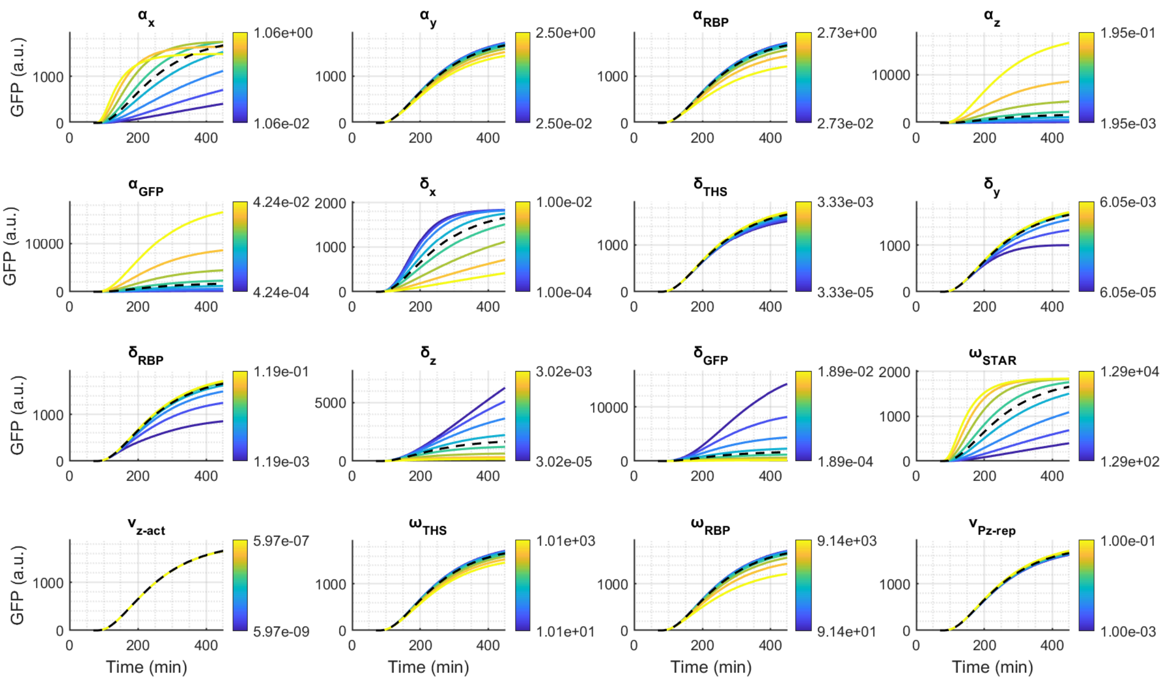


**Figure S10. Local sensitivity analysis of Circuit TL-TX-TL parameters**

Sensitivity analysis of GFP expressions to variations in 17 individual parameters while keeping all other parameters at their nominal values. Each subplot represents a different parameter, with colored lines indicating the system response across a range of parameter values (shown in the colorbar, from minimum (blue) to maximum (yellow)). The variation in GFP output (black dashed line) was analyzed by adjusting the underlying parameter values. Parameters analyzed include transcription rates (α), degradation rates (δ), binding coefficients (ω, β, γ), and decoupling coefficients (ν). Time courses demonstrate how perturbations in each parameter affect both the transient dynamics and steady-state behavior of the circuit.

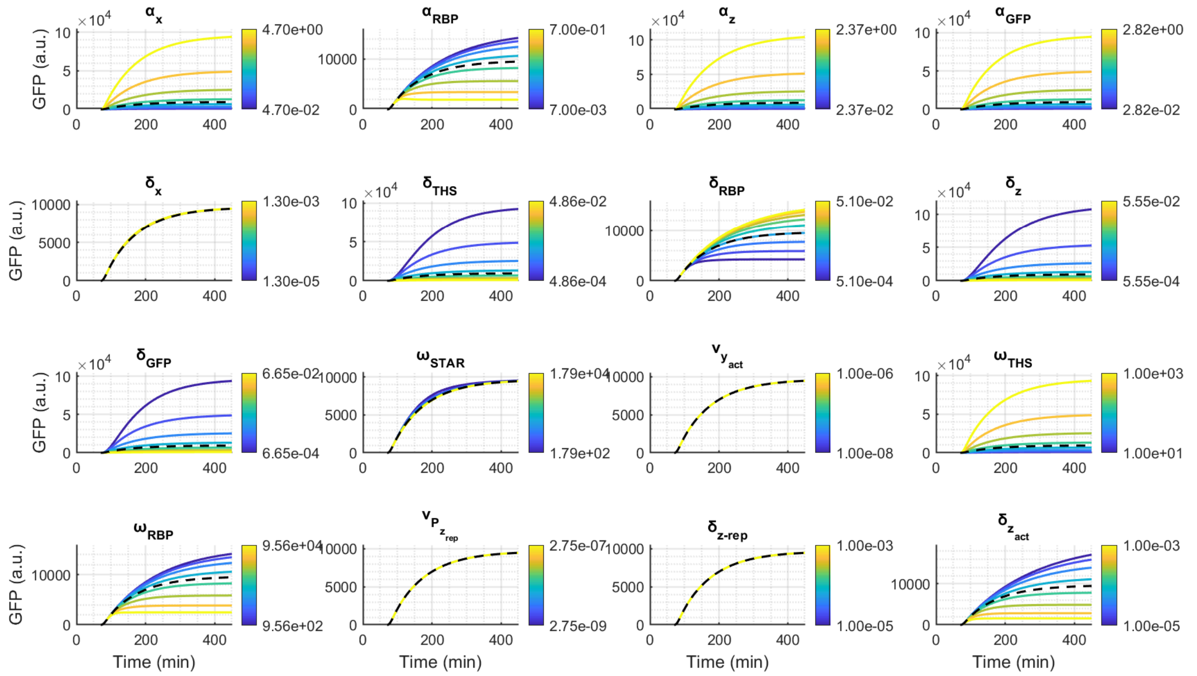


**Figure S11. Local sensitivity analysis of Circuit TX-TL-TL parameters**

The plots illustrate the system's output response (GFP expression) to variations in 16 individual parameters while keeping all other parameters at their nominal values. Each subplot represents a different parameter, with colored lines indicating the system response across a range of parameter values (shown in the colorbar, from minimum (blue) to maximum (yellow)). The variation in GFP output (black dashed line) was analyzed by adjusting the underlying parameter values. Parameters analyzed include transcription rates (α), degradation rates (δ), binding coefficients (ω, β, γ), and decoupling coefficients (ν). Time courses demonstrate how perturbations in each parameter affect both the transient dynamics and steady-state behavior of the circuit. Analysis reveals that parameters such as $\alpha_{RBP}$, $\delta_{RBP}$, $\omega_{RBP}$ strongly influence GFP expression, while others, like $\omega_{STAR}$ have minimal impact on the system's dynamics, both in transient and steady-state behaviors.

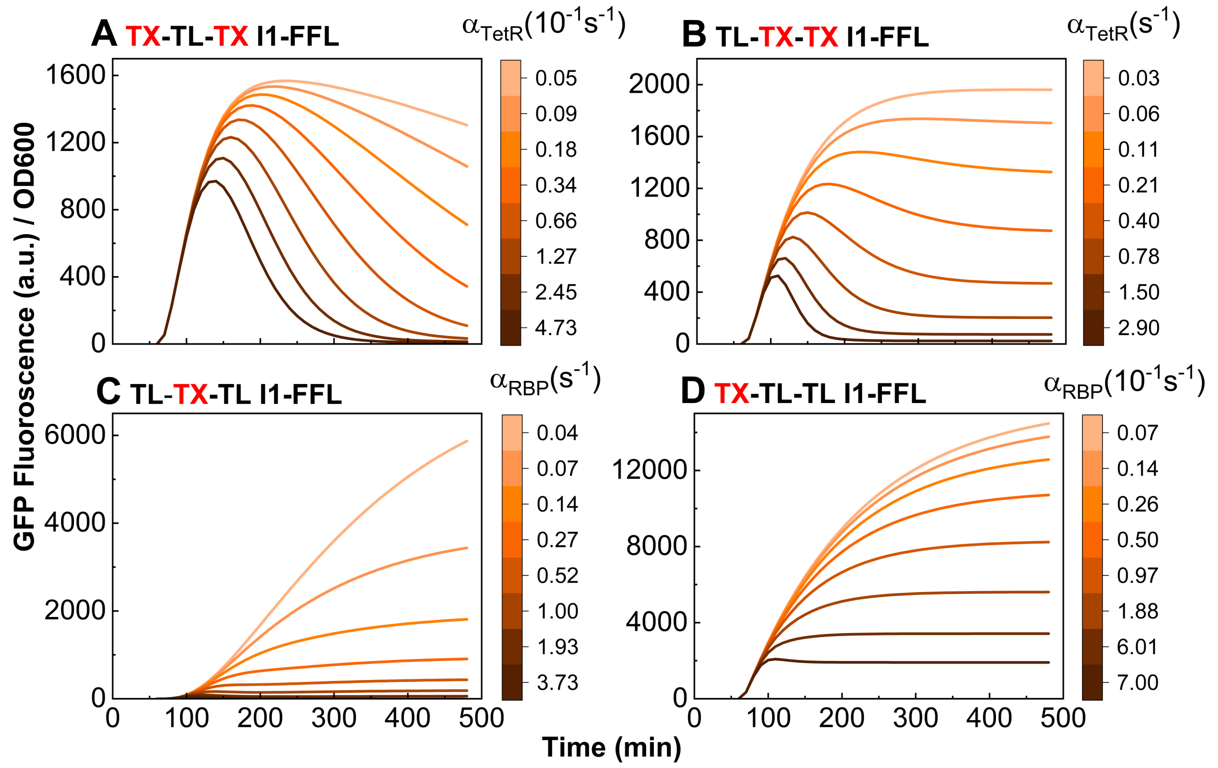


**Figure S12. Analysis of TetR/RBP Production Rate across Four I1-FFL Circuit Configurations**

The variation in GFP output was analyzed by specifically adjusting the production rate of TetR/RBP. It was varied from one-tenth of its nominal value to ten times of it in each circuit, with each range divided in 8 evenly spaced intervals to investigate its impact on circuit dynamics. The results show how changes in production rate influence the pulse generation and overall gene expression profiles for each I1-FFL circuit configuration, providing insights into the sensitivity of the circuits to variations in regulator production levels.

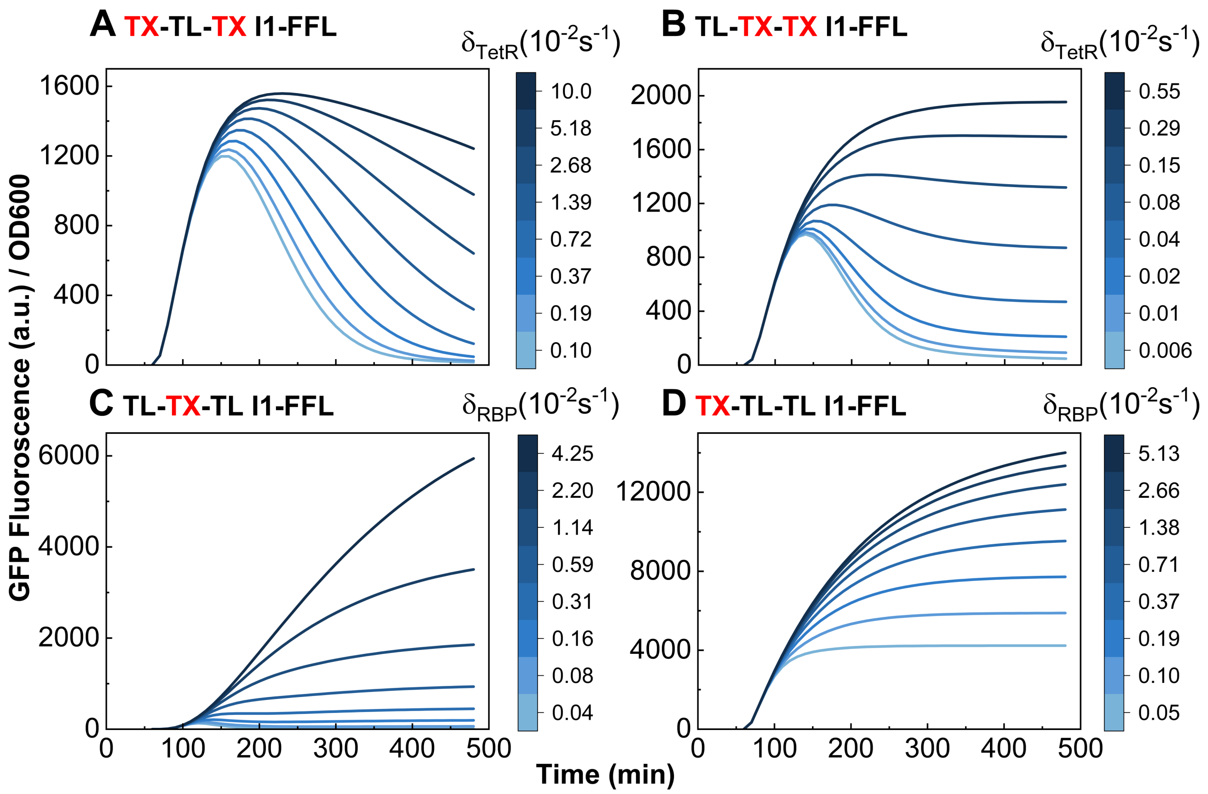


**Figure S13. Sensitivity Analysis of TetR/RBP Degradation Rate across Four I1-FFL Circuit Configurations**
The variation in GFP output was analyzed by specifically adjusting the degradation rate of TetR/RBP. It was varied from one-tenth to ten times is nominal value in each circuit, with each range divided in 8 evenly spaced intervals. The analysis reveals how degradation rate affects circuit output and stability, illustrating the importance of fine-tuning degradation for achieving the desired gene expression dynamics and maintaining pulse-like behavior in the I1-FFL circuits.

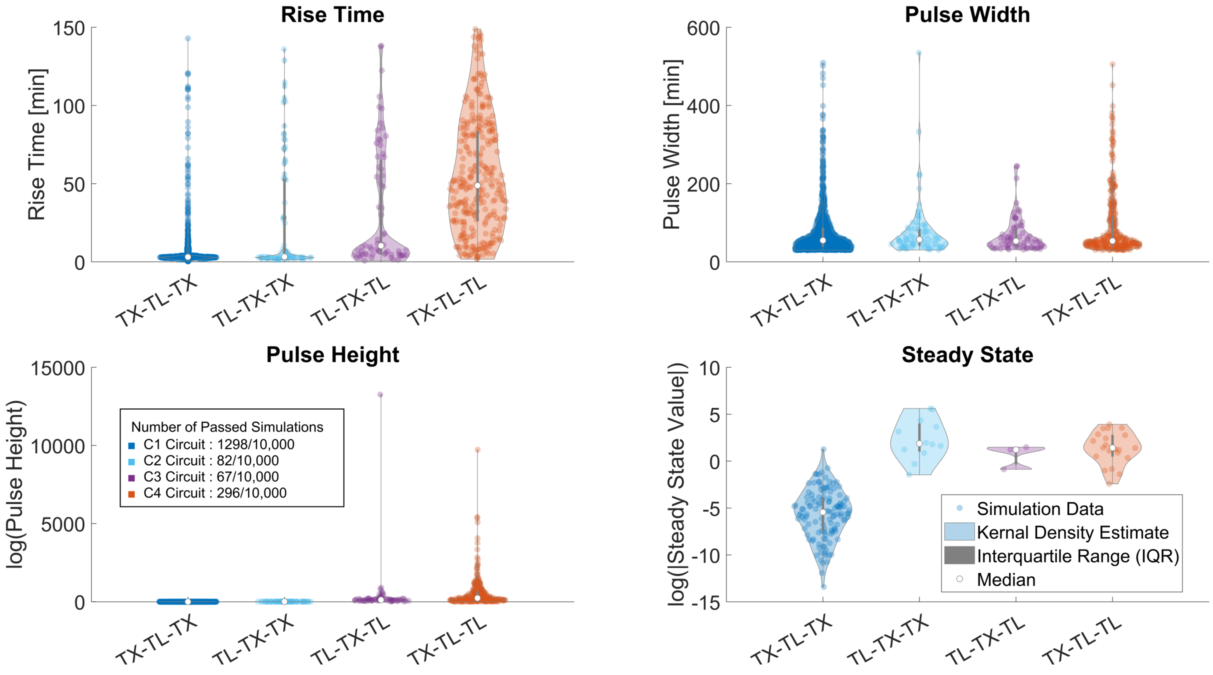


**Figure S14. Global sensitivity analysis demonstrating the dynamics achievable in each circuit**
The violin plot represents the distribution of rise time, pulse width, pulse height, and final value across the four circuits. Each point reflects the parameter value from a particular simulation; the shaded areas depict the kernel density estimation, and the dashed lines indicate the mean and quartile ranges for each distribution

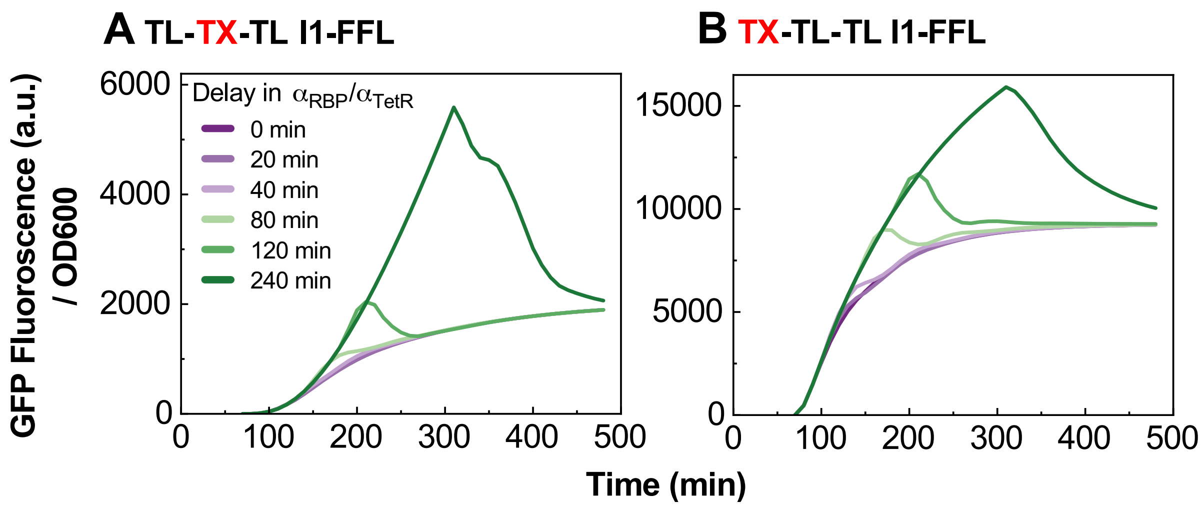


**Figure S15. Pulse generation performance in RBP circuits under simulated RBP production delays**
The graph illustrates the sharpness and timing of peaks for each circuit with varying delay periods of 20, 40, 80, 120, and 240 minutes. (a) Circuit TL-TX-TL shows peak generation at longer delays of 120 and 240 minutes. (b) whereas Circuit TX-TL-TL achieves earlier and more pronounced peaks, particularly at 80, 120, and 240 minutes.

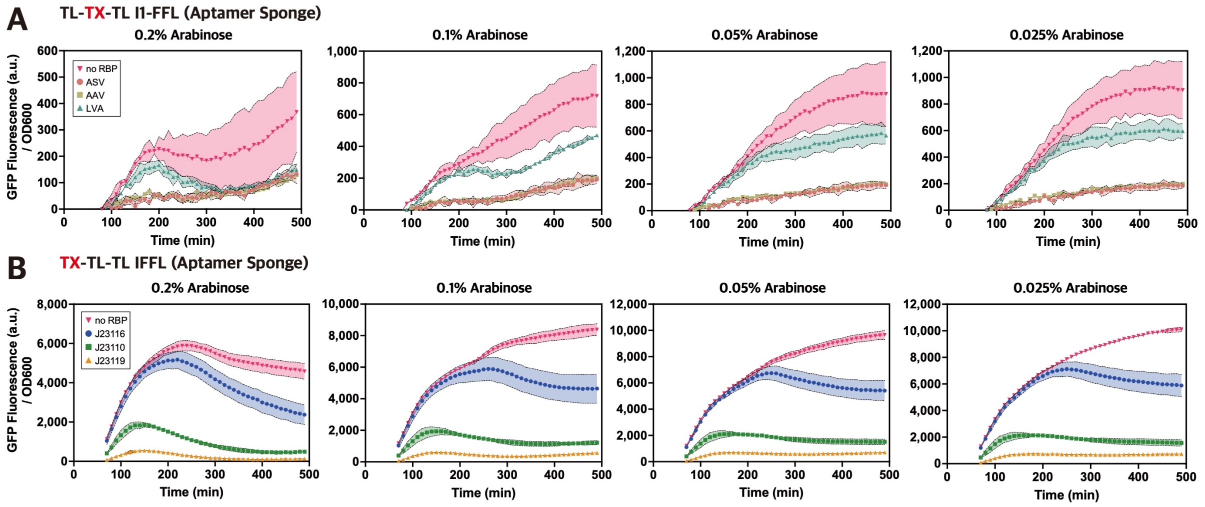


**Figure S16. Effect of Aptamer Sponge on Pulse Generation in I1-FFL circuits using RBP as a Node Y**

GFP expression dynamics in TL-TX-TL and TX-TL-TL I1-FFL circuits were monitored under four arabinose concentrations (0.2%, 0.1%, 0.05%, and 0.025%) following the introduction of an aptamer sponge to sequester PP7. All experiments measured GFP fluorescence and OD600 over a 490-minute period, with data normalized and presented as averages from three replicates. Standard deviations are shown as shaded areas. (A) TL-TX-TL I1-FFL circuit: PP7 was expressed with distinct C-terminal degradation tags to explore their impact on repression strength. Symbols represent different conditions: ASV degradation tag (salmon circles), AAV degradation tag (khaki squares), and LVA degradation tag (teal triangles). Conditions without PP7 are indicated by magenta inverted triangles. While 0.2% arabinose showed some pulse-like behavior, overall GFP expression was too low to observe distinct pulses. (B) TX-TL-TL I1-FFL circuit: Node Y was expressed under different promoters, represented as follows: J23116 (blue circles), J23110 (green squares), and J23119 (orange triangles), allowing comparisons across promoter strengths. Conditions without PP7 are indicated by magenta inverted triangles. In this circuit, pulse generation was observed under 0.2% arabinose, particularly with the J23110 and J23119 promoters, showing a clear pulse peak around 150 minutes, followed by a sharp decline in GFP levels, indicating an effective pulse response with aptamer sponge intervention.

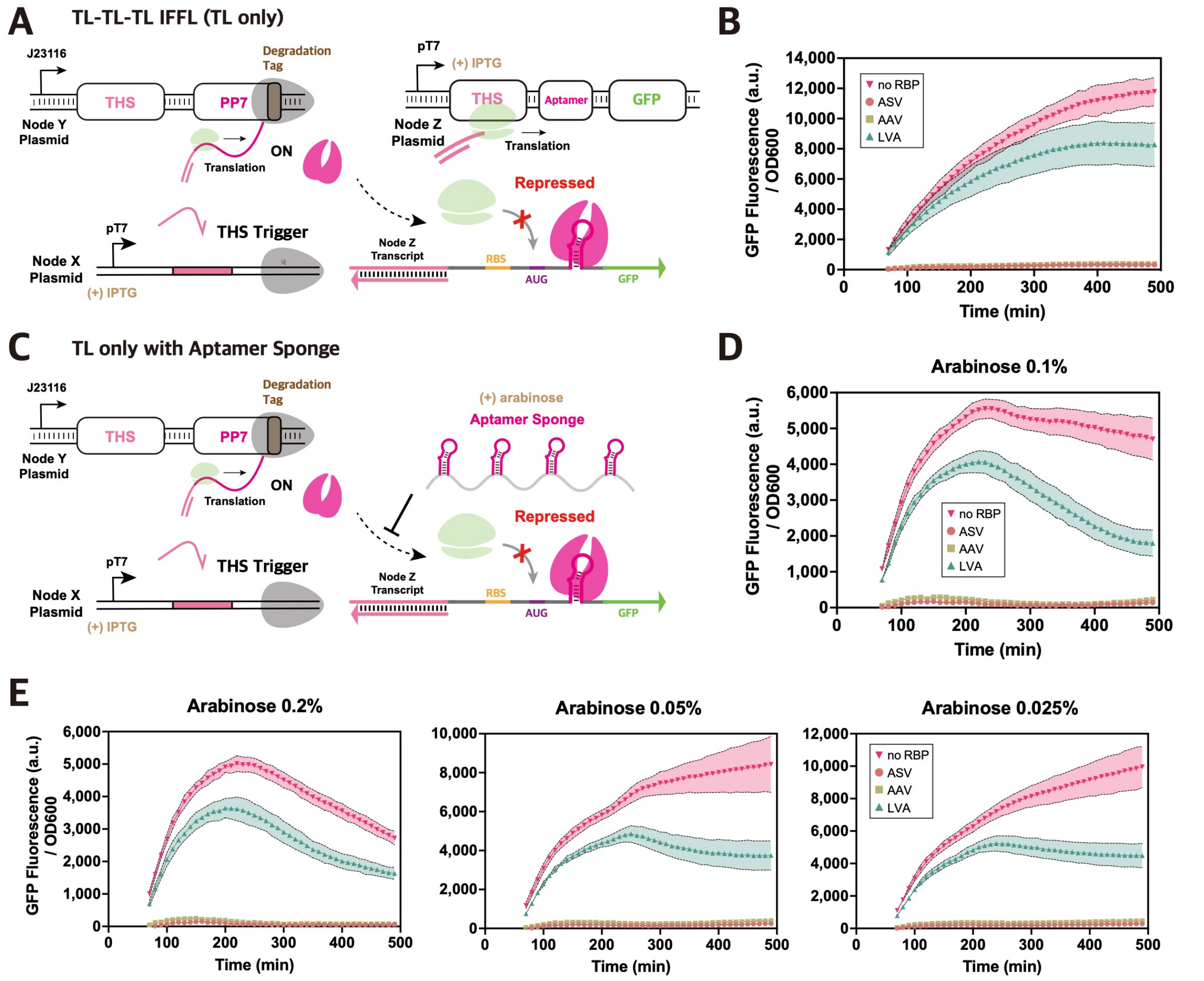


**Figure S17. Evaluation of Aptamer Sponge on Pulse Generation in TL-TL-TL(TL-only) I1-FFL Circuit**

(A) TL-only I1-FFL circuit design schematic (aptamer sponge absent): The TL-only circuit was configured with node X expressing the THS trigger, while nodes Y and Z were configured as in the TL-TX-TL and TX-TL-TL circuits, respectively. PP7, expressed via THS regulation, binds the aptamer region on node Z's transcript, blocking ribosome access and thereby inhibiting translation. (B) Experimental results without aptamer sponge: GFP expression was measured across degradation tag conditions in the TL-only circuit without an aptamer sponge. (C) TL-only I1-FFL circuit design schematic (aptamer sponge present): This setup introduced an aptamer sponge to transiently sequester PP7 early in the translation pathway. The aptamer sponge was expressed from a pBAD promoter and induced with arabinose, allowing for controlled sequestration of PP7, thereby delaying its interaction with node Z. (D) Experimental Results at 0.1% Arabinose with aptamer sponge: In the presence of 0.1% arabinose, GFP expression was measured for the TL-only circuit with aptamer sponge induction, revealing small pulse peaks under the ASV and AAV tag conditions. However, early and strong THS-driven expression of PP7 in node Y may limit the aptamer sponge's effectiveness. (e) Experimental results across varying arabinose concentrations (0.2%, 0.05%, and 0.025%): GFP expression was monitored for the TL-only circuit under different arabinose concentrations. Although pulse-like dynamics were observed, high PP7 levels at early time points continued to impact the sponge's effectiveness in generating distinct pulses. All experiments measured GFP fluorescence and OD600 over a 490-minute period, with data normalized and presented as averages from three replicates. Standard deviations are shown as shaded areas. To assess the impact of PP7 repression strength, PP7 was expressed with distinct C-terminal degradation tags: ASV (salmon circles), AAV (khaki squares), and LVA (teal triangles). Conditions lacking PP7 are indicated by magenta inverted triangles.
